# RIPK3-dependent sequential recruitment of MLKL and RIPK1 drives PANoptotic cell death and chemokine production

**DOI:** 10.1101/2025.02.26.640344

**Authors:** Yu Yang, Yue Wang, Yang Wang, Erpeng Wu, Hao Chen, Liming Sun, Chi Zhang, Shuhui Cao, Jingwen Li, Huiping Qiang, Lincheng Zhang, Yuqing Lou, Rong Qiao, Shenao Zhou, Yan Zhou, Runbo Zhong, Hua Zhong

## Abstract

PANoptosis, an immunogenic programmed cell death (PCD) modality, integrates features of pyroptosis, apoptosis, and necroptosis through assembly of the PANoptosome complex. Despite its conceptualization as a distinct PCD form, PANoptosis remains controversial due to insufficient characterization of its morphological hallmarks and molecular regulation. This study aimed to investigate the molecular mechanisms underlying the assembly of PANoptosomes in the PANoptotic pathway. We identified a novel receptor-interacting protein kinase 3 (RIPK3)-initiated PANoptotic pathway that functions without pattern recognition receptors (PRRs) and the ASC (apoptosis-associated speck-like protein containing a CARD) inflammasome. Using multimodal imaging and biochemical approaches, we identify unique morphological signatures distinguishing PANoptotic cells from canonical pyroptotic, apoptotic, or necroptotic counterparts. Mechanistically, RIPK3 forms round homopolymeric scaffolds—distinct from necroptotic amyloid-like fibers—to sequentially recruit MLKL and RIPK1, forming a dynamic RIPK3-MLKL-RIPK1-FADD-caspase-8 complex (RIPK3-PANoptosome). This platform coordinates concurrent activation of pyroptotic, apoptotic, and necroptotic effectors. Cross-regulatory interactions between these pathways establish a homeostatic system where perturbations bias death modality into a certain cell death type, altering the death process and outcomes. Functionally, PANoptotic cells orchestrate chemokine secretion through parallel kinase-dependent (RIPK3-MLKL) and kinase-independent (RIPK1-IKK-NF-κB) mechanisms, driving macrophage recruitment. Our findings resolve the molecular logic of PANoptosome assembly, redefine PANoptosis as a tunable PCD paradigm, and establish its role in immunomodulation, providing a framework for targeting inflammatory cell death in disease.

## Introduction

Programmed cell death (PCD) is a process by which cells actively terminate their lives and is crucial in maintaining tissue homeostasis, regulating organismal development, and defending against pathogen invasion. Receptor-interacting protein kinase 3 (RIPK3) is a vital regulator of various PCD pathways^1–3^. RIPK3 is activated in response to multiple cellular stress stimuli via its corresponding death receptors (DRs), such as TNFR^4–6^, DR4/5^7^, and DR6^8^, pattern recognition receptors, such as TLR3/4^9,10^ and ZBP1^11–17^, or viral protein ICP6^18^. This activation induces the formation of large polymers within different protein complexes. Once activated via oligomerization and autophosphorylation, RIPK3 is primed to recruit mixed lineage kinase domain-like (MLKL) proteins from the cytosol, phosphorylate the activation loop within the MLKL C-terminal domain, and facilitate the plasma membrane translocation of MLKL, triggering necroptosis^19–23^. Under certain conditions, such as the ectopic expression of the D161N mutant^24,25^, influenza A virus (IAV) infection^13,26,27^, treatment with small-molecule inhibitors^24^, or the inhibition of the Hsp90/CDC37 chaperone^28^, RIPK3 promotes apoptosis via the RIPK1-FADD-Caspase-8 pathway. In addition, RIPK3 plays a regulatory role in inflammasome assembly^29^, triggering pyroptotic cell death, highlighting the significance of RIPK3 in immune responses and inflammatory diseases.

A new PCD pathway that integrates pyroptosis, apoptosis, and necroptosis features, termed PANoptosis, was recently characterized^30–35^. This pathway is driven by PANoptosome complexes, which comprise innate immune sensors (including ZBP1, AIM2, NLRP3, and NLRP12), adaptors (including ASC and FADD), catalytic effectors (including caspase-1, −6, −8, RIPK1, and RIPK3), and executors (including MLKL)^36^. Four PANoptosome complexes (ZBP1-^30,31^, AIM2-^32^, RIPK1-^33^, and NLRP12-^34,35^PANoptosomes) have been identified, and their assembly occurs in response to pathogen-associated molecular patterns (PAMPs) derived from microbial infections^30–32^ and damage-associated molecular patterns (DAMPs) from damaged or dying cells^34,35^. However, the events downstream of PANoptosis-associated sensor activation remain unclear. The comprehensive mechanisms underlying PANoptosis, such as PANoptosome assembly and the interplay between different PCD pathways, are partially understood. This has prompted controversy regarding whether PANoptosis should be identified as a separate PCD type or merely as an amalgamation of the existing forms.

This study aimed to investigate the molecular mechanisms underlying the assembly of PANoptosome in the PANoptotic pathway, a novel PCD form. We discovered a novel RIPK3-initiated PANoptotic pathway without PPRs and the ASC complex and elucidated the molecular mechanisms underlying the assembly of the associated complexes. We demonstrated that the direct activation of RIPK3, such as via osmotic stress (OS) stimulation, chemically inducible dimerization of RIPK3, or artificial ectopic overexpression of RIPK3 or RIPK1, induces the formation of RIPK3-MLKL-RIPK1-FADD-Caspase-8-PANoptosome complexes, triggering RIPK3-PANoptosis and mediating chemokine secretion. We observed phospho-MLKL, marked for necroptosis, and cleaved Caspase-8, as upstream proteases for Caspase-3 and gasdermin D (GSDMD) processing in a single cell, where PANoptosomes at different stages were also detected. Our results suggest that the observed activation of different PCD pathways during RIPK3-PANoptosis was induced by an integrated system formed by interacting pathways rather than by a mix of distinct cells that underwent different PCDs. These molecular events that occurred in a single cell facilitated PANoptotic cells to exhibit unique morphological features and cytokine secretion patterns. Our findings provide new insights into PANoptosis as a distinct cell death form.

## Results

### Direct RIPK3 activation orchestrates PANoptotic cell death

The restored RIPK3 expression potentiated the sensitivity of PKLCs to OS and caused cell death (Fig. 1a). OS treatment triggered MLKL phosphorylation, as previously reported^37^, and induced caspase and GSDMD cleavage (Fig. 1b). Similar results were observed in the RIPK3-restored human PL45 cells (Supplementary Fig. 1a). To clarify the contribution of these death signals to OS-induced cell death, we evaluated OS-triggered cell loss following pretreatment with the RIPK3 kinase inhibitor GSK’872, pan-caspase inhibitor Z-VAD, or both using crystal violet staining. Only the combined GSK’872 and Z-VAD treatment significantly inhibited OS-induced cell death, while GSK’872 or Z-VAD alone did not (Fig. 1c). Immunoblotting showed that GSK’872 and Z-VAD respectively blocked the MLKL phosphorylation and Caspase-8, Capspase-3, and GSDMD cleavage in the OS-stimulated cells, and GSK’872 plus Z-VAD inhibited MLKL phosphorylation compared with the effect of treatment with Z-VAD alone and completely blocked caspase and GSDMD cleavage (Fig. 1d). Thus, we confirmed that phospho-MLKL and cleaved caspases contributed to OS-induced cell death.

**Fig. 1.**
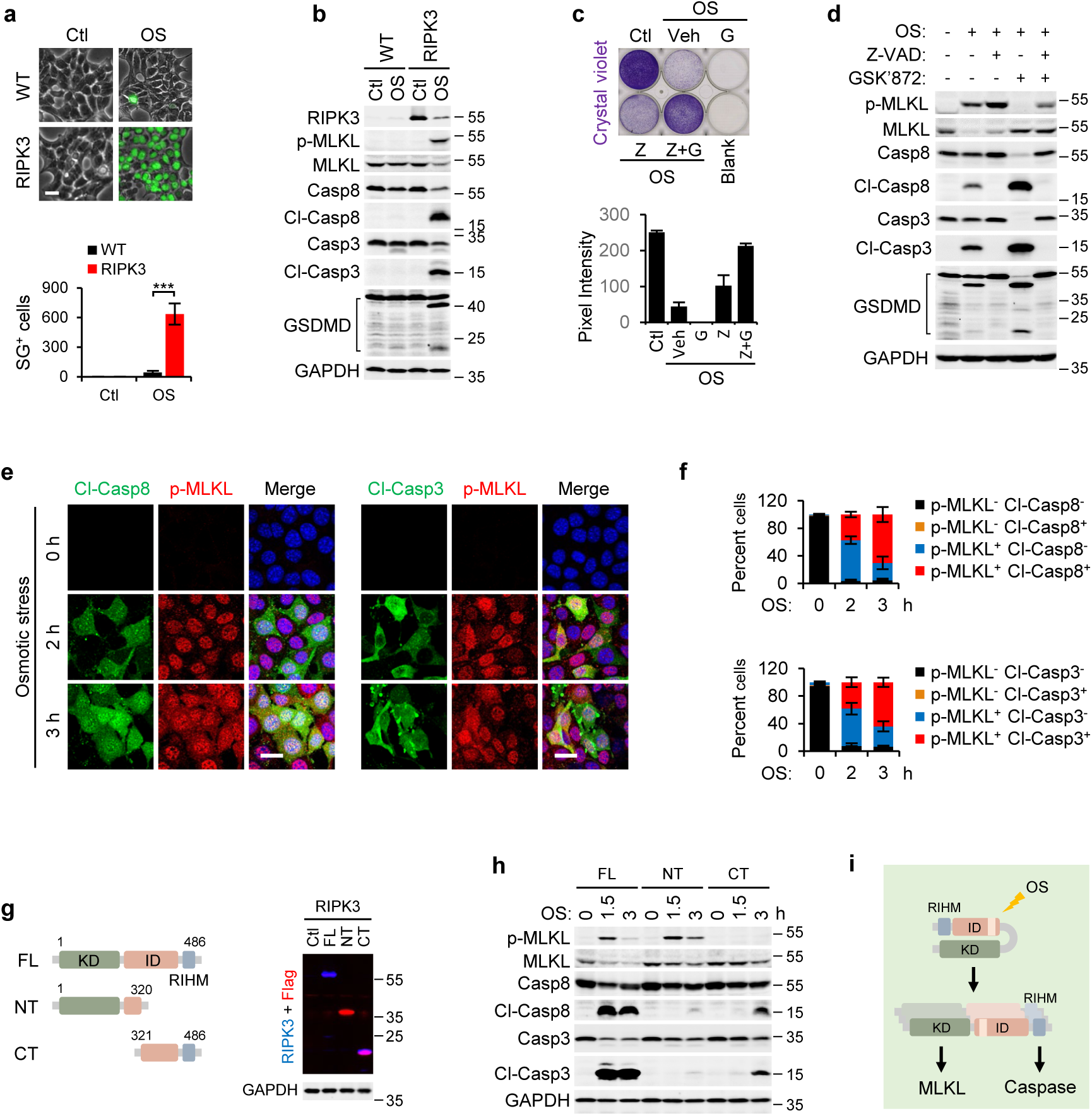
Osmotic stress drives RIPK3-mediated MLKL phosphorylation and caspase activation concurrently in single cells. **a** Wild-type (WT) or RIPK3-restored PKLCs were treated with PBS (control, Ctl) or 400 mM NaCl (osmotic stress, OS) for 4 hours. Plasma membrane permeabilization was assessed using the SYTOX Green (SG) uptake assay. SG-positive cells were quantified using ImageJ. Representative images from three independent experiments are shown. Scale bar: 20 μm. **b** WT or RIPK3-restored PKLCs were subjected to OS for 1.5 hours. Levels of RIPK3, p-MLKL (necroptosis markers), cleaved Caspase-8 (Cl-Casp8), cleaved Caspase-3 (Cl-Casp3) (apoptosis markers), and GSDMD (pyroptosis executor) in cell lysates were analyzed by immunoblotting. GAPDH served as the loading control. Blots are representative of three independent experiments. **c, d** RIPK3-restored PKLCs were treated with OS alone or in combination with Vehicle (Veh), Z-VAD (Z), GSK’872 (G), or both (Z+G) for 2 hours. Culture media were replaced with fresh media, and cells were incubated for an additional 12 hours. Cell viability was determined using the crystal violet assay. Representative images from three independent experiments are shown. **e, f** Representative immunofluorescence images (**e**) and quantification (**f**) showing the percentage of RIPK3-restored PKLC cells with distinct signals after OS treatment for the indicated durations. Cells were stained with antibodies against p-MLKL and Cl-Casp8 or p-MLKL and Cl-Casp3. Scale bar: 20 μm. Data are presented as means ± SD from eight fields. g, h PKLC cells were stably reconstituted with full-length (FL), N-terminal-truncated (NT, 1-320 aa), or C-terminal-truncated (CT, 321-486 aa) forms of RIPK3 (**g**) and treated with OS for the indicated durations (**h**). Whole-cell lysates were analyzed by immunoblotting using the indicated antibodies. **i** Schematic illustrating RIPK3-dependent MLKL phosphorylation and caspase cleavage induced by OS.

ZBP1 reportedly activates the parallel pathways of MLKL-mediated necroptosis and caspase-mediated apoptosis by sensing the genomic RNA of IAV, evidenced by MLKL phosphorylation and caspase cleavage, as revealed by immunoblotting after IAV infection^13,26^. However, these two cell death pathways are not activated in the same infected cell and restrict IAV infection independently^27^. To investigate whether OS-induced cell death has a similar mechanism, we performed co-immunostaining for phospho-MLKL with cleaved Caspase-8 or −3 in RIPK3-restored PKLCs treated with OS. MLKL phosphorylation and caspase cleavage were observed in the same cell (Fig. 1e). The proportion of cells double-positive for phospho-MLKL and cleaved Caspase-8 gradually increased to approximately 70% over 3 h, and that of cells that were positive for phospho-MLKL and cleaved Caspase-3 was largely equivalent (Fig. 1f).

In OS-induced necroptosis, NHE1 triggers an increase in the intracellular pH to activate the RIPK3 N-terminal kinase domain, causing MLKL phosphorylation^37^. To test whether RIPK3-initiated cleavage of caspase and GSDMD proceeded via a similar mechanism, we introduced N- or C-terminally truncated RIPK3 into PKLCs (Fig. 1g). As expected, MLKL was phosphorylated, and negligible caspase was processed in response to OS in cells expressing N-terminally truncated RIPK3^37^. In contrast, cleaved caspases, but not phospho-MLKL, were observed in OS-stimulated cells expressing C-terminally truncated RIPK3 (Fig. 1h). Hyperosmotic stress directly induced a conformational switch in RIPK3, causing the separate activation of its N-terminal kinase and C-terminal RIHM domains to mediate MLKL phosphorylation and caspase cleavage (Fig. 1i).

Hyperosmotic stress activates numerous processes that remain largely uncharacterized^38^. With prolonged OS exposure, the cells died in an RIPK3-independent manner. To investigate the events downstream of RIPK3 activation, we engineered RIPK3^2×Fv^-expressing and doxycycline (Dox)-inducible RIPK3-expressing PKLC cell lines. The former expressed RIPK3 fused with a Flag tag and the tandem-binding domain of FKBP12 (Fv) at the C-terminus, allowing the rapid activation of RIPK3 by the addition of a B/B homodimerizer. Consistent with OS-induced RIPK3 activation, RIPK3 oligomerization in PKLCs caused robust cell death, which was blocked by the combination treatment with Z-VAD and GSK’872 but not by either reagent alone (Fig. 2a, b). To verify this hypothesis, we treated PKLC-RIPK3^2×Fv^ cells with a dimerizer (Dim) and analyzed the biochemical markers of necroptosis, apoptosis, and pyroptosis at different time points. A strong phospho-MLKL signal appeared at an early stage after treatment and continued to increase; however, it was inhibited by treatment with the RIPK3 kinase inhibitor GSK’872 (Fig. 2c, d). Moreover, the cleavage of apoptotic markers, such as Caspase-8, Caspase-3, and their substrate PARP, occurred 1 h after treatment, later than the MLKL phosphorylation. In addition, the pyroptosis executor GSDMD was cleaved into its p40, p30, and p20 fragments simultaneously with caspase processing (Fig. 2c). The auto-phosphorylation of RIPK3 at Ser165 and Thr166 is required to initiate intracellular apoptosis through the RIPK1-FADD-Caspase-8 pathway, which depends on RIPK3 kinase activity^28^. Conversely, the cleavage events in our study could not be blocked by treatment with RIPK3 kinase inhibitor GSK’872 (Fig. 2d). In PKLC-RIPK3^Tet-on^ cells, the ectopic expression of RIPK3 is controlled by a Tet operator and induced by Dox. Dox-induced cell death began 2 h after Dox treatment and reached a plateau after 12 h (Fig. 2e). Similar to RIPK3 oligomerization, RIPK3 overexpression-triggered cell death was blocked by the combination treatment with Z-VAD and GSK’872 but not by Z-VAD or GSK’872 alone (Fig. 2f). Corresponding to cell death, RIPK3 was expressed 2 h after Dox stimulation, and its expression was increased over time and decreased 4 h after treatment (Fig. 2g). As expected, we detected the simultaneous activation of MLKL and cleaved caspases, which were blocked by GSK’872 and Z-VAD, respectively (Fig. 2g, h). The contemporaneous activation of MLKL and caspases in response to RIPK3 was confirmed using two RIPK3^2×Fv^-restored human cell lines, PL45-RIPK3^2×Fv^ and 786-O-RIPK3^2×Fv^. RIPK3 oligomerization-induced cell death in PL45 and 786-O cells was slightly attenuated by the necroptosis inhibitor necrosulfonamide (NSA) and blocked by the combination treatment with NSA and Z-VAD (Supplementary Fig. 1b, c). Phospho-MLKL and cleaved-PARP were detected in PL45 and 786-O cells 3 and 6 h, respectively, after treatment (Supplementary Fig. 1d, e). Because of the low gasdermins expression, we did not observe GSDMD or GSDME cleavage in PL45 cells (Supplementary Fig. 1d). In 786-O cells, GSDME was cleaved into its p32 fragment (Supplementary Fig. 1e). Previous studies have shown that RIPK3 dimerization primarily causes MLKL-mediated necroptosis. Thus, we examined RIPK3-initiated cell death in Lewis lung carcinoma (LLC) and NIH-3T3 cells. The cell death in both cell types was completely blocked by GSK’872 (Supplementary Fig. 1f, g), and a cleaved caspase or GSDMD signal was not detected during the cell death (Supplementary Fig. 1h, i). These data indicate that LLC and NIH-3T3 cells underwent necroptotic cell death induced by RIPK3 oligomerization. Hence, we presume that RIPK3 induces caspase activation in a cell type-specific manner via unknown mechanisms.

**Fig. 2.**
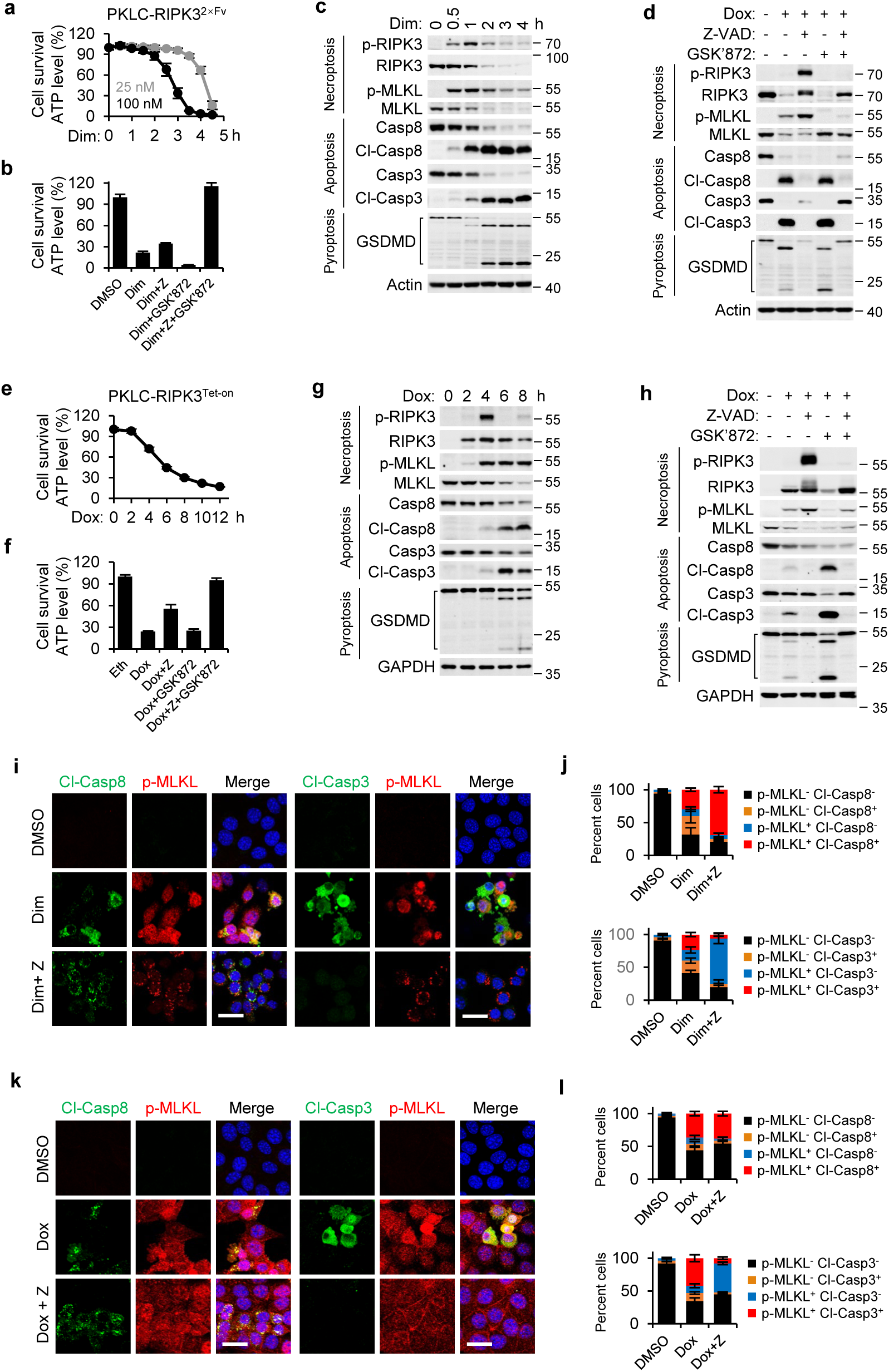
Direct RIPK3 activation orchestrates PANoptotic cell death. **a-d** PKLC-RIPK3^2×Fv^ cells were treated with 25 nM or 100 nM Dim for the indicated durations (**a, c**), or with Dim, Dim + Z-VAD (Z), Dim + GSK’872, Dim + Z + GSK’872 for 4 hours (**b, d**). Cell viability was determined by measuring ATP levels using the CellTiter-Glo kit (**a, b**). Data are presented as means ± SD of triplicate wells. Levels of p-RIPK3, p-MLKL (necroptosis markers), cleaved Caspase-8 (Cl-Casp8), cleaved Caspase-3 (Cl-Casp3) (apoptosis markers), and cleaved GSDMD (Cl-GSDMD, pyroptosis marker) were detected by immunoblotting (**c, d**). Results are representative of three independent experiments. **e-h** PKLC-RIPK3^Tet-on^ cells were treated with Dox for the indicated durations (**e, g**), or with Dox, Dox + Z, Dox + GSK’872, Dox + Z + GSK’872 for 8 hours (**f, h**). Cell viability was determined by measuring ATP levels using the CellTiter-Glo kit (**e, f**). Data are presented as means ± SD of triplicate wells. Cell lysates were analyzed by immunoblotting using the specified antibodies (**g, h**). Results are representative of three independent experiments. **i-l** Representative immunofluorescence images (**i, k**) and quantification showing the percentage of cells with distinct signals (**j, l**) in PKLC-RIPK3^2×Fv^ cells treated with DMSO, Dim, or Dim + Z for 2 hours (**i, j**), and PKLC-RIPK3^Tet-on^ cells treated with DMSO, Dox, or Dox + Z for 6 hours (**k, l**). Cells were stained with antibodies against p-MLKL and Cl-Casp8 or p-MLKL and Cl-Casp3. Scale bar, 20 μm. Data are presented as means ± SD of eight fields.

To characterize the activation of death signals at the single-cell level, we performed co-immunostaining for phospho-MLKL with cleaved Caspase-8 or −3 in PKLC-RIPK3^2×Fv^ cells treated with vehicle (dimethyl sulfoxide, DMSO), Dim, or Dim plus Z-VAD (DZ). Consistent with immunoblotting, MLKL phosphorylation was observed in approximately 40% of dimerizer-treated cells, and this proportion increased to 70% in cells treated with DZ (Fig. 2i, j). We previously found that Z-VAD did not affect the primary cleavage of pro-Caspase-8 at Asp373 and Asp387, resulting in the generation of p41/p43 and p10 fragments; however, it completely inhibited further cleavage of Caspase-8 at Asp212 and Asp218, which released the large catalytic p18 fragment. Accordingly, the immunofluorescence signal of cleaved Caspase-8 was observed in Dim-treated cells with or without Z-VAD (Fig. 2i, j). Phospho-MLKL/cleaved Caspase-8-double-positive cells were also observed (Fig. 2i). As the suppression of the catalytic activity of caspases blocked the dissociation of the caspase-associated complex, leading to the accumulation of upstream signals, an increased percentage, from approximately 30% to 70%, of phospho-MLKL/cleaved Caspase-8 double-positive cells was observed following treatment with the DZ, compared with the effect of treatment with the Dim alone (Fig. 2j). Cleaved Caspase-3 was detected in phospho-MLKL-positive cells and was strongly inhibited by Z-VAD (Fig. 2i). Statistical analysis showed that over 23% of the cells were phospho-MLKL/cleaved Caspase-3-double positive 2 h after treatment (Fig. 2j). Although the proportion of phospho-MLKL/cleaved caspase-double positive PKLC-RIPK3^Tet-on^ cells treated with Dox or Dox plus Z-VAD was relatively small owing to the rapid detachment of dying cells, the death signals displayed similar patterns to those in PKLC-RIPK3^2×Fv^ cells treated with the Dim or DZ (Fig. 2k, l).

Despite the limited commercially available antibodies against cleaved GSDMD for immunofluorescence, GSDMD was completely cleaved 2 h after stimulation, as revealed by immunoblotting with an antibody against total GSDMD (Fig. 2c). Hence, we hypothesized that MLKL phosphorylation, caspase cleavage, and GSDMD cleavage occurred simultaneously within a single cell in response to RIPK3 oligomerization. Thus, we demonstrated that direct RIPK3 activation could initiate PANoptotic cell death, termed RIPK3-PANoptosis.

### RIPK scaffolding mediates caspase activation during RIPK3-PANoptosis

To identify the molecules downstream of RIPK3, Flag-tagged RIPK3 was isolated from DMSO- or Dim-treated PKLC-RIPK3^2×Fv^ cells using anti-Flag beads and subjected to mass spectrometry. RIPK1 and Caspase-8 coprecipitated with RIPK3 upon dimerization (data not shown), indicating that these molecules interact with RIPK3 during cell death. These results suggest that the observed caspase cleavage was mediated by the RIPK1-FADD-Caspase-8 pathway. Therefore, we depleted RIPK1 in wild-type (WT) and MLKL-deficient PKLC-RIPK3^2×Fv^ cells using shRNA and found that, in contrast to TNF plus Smac mimetic and Z-VAD (TSZ)-induced necroptosis, RIPK1 knockdown failed to block Dim-induced cell death in MLKL-expressing cells (Fig. 3a). Immunoblotting revealed that RIPK1 knockdown did not affect RIPK3 and MLKL phosphorylation but completely blocked Caspase-8 and −3 cleavage (Fig. 3b). Correspondingly, RIPK1 deficiency restricted Dim-induced cell death in MLKL-deficient cells and blocked the cleavage of Caspase-8, Caspase-3, and GSDMD (Fig. 3c, d). Furthermore, RIPK1 deficiency suppressed the interaction between Caspase-8 and RIPK3 in MLKL-expressing and -deficient cells (Supplementary Fig. 2a, b). Subsequently, we depleted FADD and Caspase-8 in MLKL-deficient PKLC-RIPK3^2×Fv^ cells. As expected, FADD or RIPK1 knockdown blocked the Dim-induced apoptosis in MLKL-deficient cells (Fig. 3e, f). These indicate that RIPK1 is crucial in the assembly of the RIPK3-RIPK1-Caspase-8 complex and caspase activation during RIPK3-PANoptosis.

**Fig. 3.**
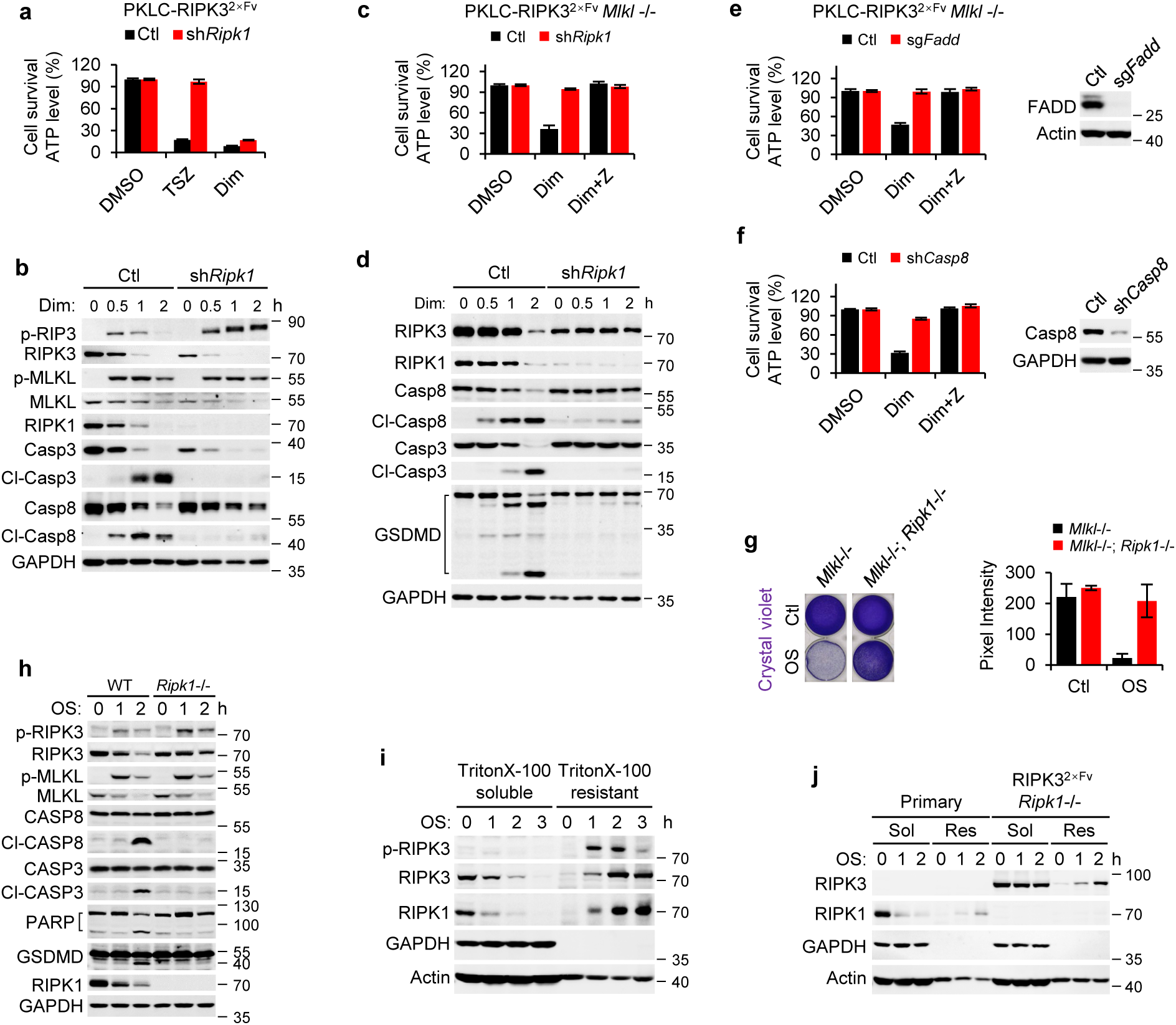
RIPK1–FADD axis drives caspase processing downstream of RIPK3 during RIPK3-PANoptosis. **a, b** WT or RIPK1-deficient PKLC-RIPK3^2×Fv^ cells were treated with DMSO, TSZ for 6 hours, or Dim for 4 hours (**a**), or for the indicated durations (**b**). Cell viability was determined by measuring ATP levels using the CellTiter-Glo kit (**a**). Data are presented as means ± SD of triplicate wells. Whole-cell lysates were analyzed by immunoblotting using the specified antibodies (**b**). **c, d** MLKL knockout PKLC-RIPK3^2×Fv^ cells with WT or RIPK1-deficient genotypes were treated with DMSO, Dim, or Dim + Z for 4 hours (**c**), or for the indicated durations (**d**). Cell viability was determined by measuring ATP levels using the CellTiter-Glo kit (**c**). Data are presented as means ± SD of triplicate wells. Whole-cell lysates were analyzed by immunoblotting using the specified antibodies (**d**). **g, h** MLKL knockout PKLC-RIPK3^2×Fv^ cells with WT or RIPK1-deficient genotypes were treated with PBS or OS for 1 hour (**g**), or for the indicated durations (**h**). Cell viability was determined by crystal violet assay (**g**). Data are presented as means ± SD of triplicate wells. Whole-cell lysates were analyzed by immunoblotting using the specified antibodies (**h**). **i, j** Immunoblotting analysis of p-RIPK3 (**i**), RIPK3, and RIPK1 (**i, j**) localization in Triton X-100-soluble and Triton X-100-resistant fractions from primary PKLC, RIPK1-deficient PKLC-RIPK3^2×Fv^ (**j**), or PKLC-RIPK3^2×Fv^ (**i**) cells treated with OS for the indicated durations.

To determine whether OS-induced caspase activation also depends on RIPK1, we examined cell loss via the crystal violet staining of MLKL-deficient or MLKL-/RIPK1-double deficient PKLC-RIPK3^2×Fv^ cells. MLKL-deficient parental PKLC-RIPK3^2×Fv^ cells died after OS exposure for 1 h, resulting in weak staining in the culture plates. MLKL-/RIPK1-double deficient PKLC-RIPK3^2×Fv^ cells survived the same OS condition, with indistinguishable stained culture plates from those of the control group (Fig. 3g). Consistently, RIPK1 deficiency blocked Caspase-8/-3, PARP, and GSDMD cleavage but did not affect RIPK3 or MLKL phosphorylation, indicating that the OS-induced caspase activation depended on RIPK1 (Fig. 3h). Similarly, RIPK3 and RIPK1 translocated to the TritonX-100 resistant fraction after OS induction, suggesting an RIHM domain-mediated polymerization of these RIPKs in response to OS stimuli (Fig. 3i). Furthermore, a significantly decreased translocation of RIPK1 to the TritonX-100 resistant fraction occurred in OS-treated WT PKLCs that did not express RIPK3, whereas RIPK3 translocation was not affected in the OS-treated RIPK1-deficient PKLC-RIPK3^2×Fv^ cells (Fig. 3j). These data show that RIPK1 acted downstream of RIPK3 in OS-exposed cells.

RIPK1 contains an N-terminal kinase domain, a C-terminal death domain, and an intermediate domain with an RHIM motif^39^. While RIPK1 kinase activity is crucial for RIPK1-RIPK3-MLKL necrosome formation to induce necroptosis^40,41^, the RIPK1 kinase inhibitor Nec-1s did not attenuate RIPK3 dimerization-induced cell death in the MLKL-expressing or -deficient PKLCs (Supplementary Fig. 3a, b). Immunoblotting showed that MLKL phosphorylation and Caspase-8/-3 and GSDMD cleavage were not affected by Nec-1s (Supplementary Fig. 3c, d), suggesting that PANoptotic death is not dependent on RIPK1 kinase activity. The ectopic expression of RIPK1^D325∼end^, a C-terminal truncation lacking the kinase domain, rescued caspase activation and apoptosis in MLKL/RIPK1-double deficient PKLC-RIPK3^2×Fv^ cells (Supplementary Fig. 3e, f).

We previously found that GSK’872 alone could not inhibit PANoptosis but completely inhibited TSZ-induced necroptosis at the same concentration (Supplementary Fig. 3g). Additionally, PANoptosis was blocked by the combination treatment with Z-VAD and GSK’872, indicating that the inhibition of RIPK3 kinase activity switches PANoptosis to caspase-dependent cell death. Following treatment with Dim plus GSK’872 (DG), the cells exhibited typical apoptotic morphology, such as an intact plasma membrane, nuclear condensation, and apoptotic body formation (Supplementary Fig. 3h). The dynamics of cell death in DG-induced apoptosis were markedly similar to those in Dim-induced PANoptosis and different from those of Dim-induced apoptosis in MLKL-deficient cells, suggesting that GSK’872 exacerbated RIPK3 dimerization-induced apoptosis (Supplementary Fig. 3i). To confirm this, we compared Dim-induced apoptosis with or without GSK’872 treatment in MLKL-deficient cells. GSK’872 significantly enhanced caspase-dependent cell death in a dose-dependent manner (Supplementary Fig. 3j). Immunoblotting revealed strongly enhanced signals for cleaved Caspase-8, Caspase-3, and GSDMD in PANoptotic cells after the combination treatment with GSK’872 (Supplementary Fig. 3k). To explain the effect of RIPK3 kinase activity on caspase activation, we performed immunoprecipitation of Flag-tagged RIPK3 from PANoptotic cells with or without GSK’872 treatment. The inhibition of RIPK3 kinase activity increased its affinity for RIPK1 and facilitated the assembly of the RIPK3-RIPK1-Caspase-8 complex (Supplementary Fig. 3l). Similar to RIPK1, RIPK3 comprises an N-terminal kinase domain and a C-terminal RHIM motif. We engineered a Dox-inducible RIPK3^D333∼end^-expressing PKLC cell line and found that the ectopic expression of the RIPK3^D333∼end^ was sufficient to mediate caspase processing and caspase-dependent but not MLKL-dependent cell death, which was inhibited by Z-VAD (Supplementary Fig. 3m, n). These results indicate that the scaffold function, and not the kinase activity of RIPK1 and RIPK3, is essential for caspase activation during RIPK3-PANoptosis. Although RIPK3 kinase activity is crucial for MLKL phosphorylation, it negatively regulates caspase activation.

### The affinity for RIPK3 coordinates stepwise assembly of MLKL and RIPK1 into the PANoptosome

We previously observed that RIPK3 recruits MLKL, RIPK1, and Caspase-8 to form a multimeric protein complex upon dimerization (Supplementary Fig. 2a, b, Fig. 4a). This RIPK3-MLKL-RIPK1 complex formation was confirmed through Flag-RIPK3 immunoprecipitation in PKLC-RIPK3^Tet-on^ (Fig. 4b) and PL45-RIPK3^2×Fv^ (Fig. 4c) cells. The interaction between RIPK1 and RIPK3 continued to increase during cell death, while that between MLKL and RIPK3 initially increased and subsequently declined (Fig. 4a-c). To investigate the molecular mechanisms of these complex assemblies, we immunoprecipitated Flag-RIPK3 and V5-MLKL from Dim- and DZ-treated PKLC-RIPK3^2×Fv^ cells and analyzed their molecular compositions via immunoblotting. The assay was performed using N-terminal V5-tagged MLKL-expressing cells to eliminate the effects of lytic cell death, which causes a rapid loss of cell content. RIPK1 and Caspase-8 were recruited to RIPK3 upon Dim treatment, which was augmented by Z-VAD owing to the blockage of downstream events (Fig. 4d). As observed in non-tagged MLKL-expressing cells, MLKL bound to RIPK3 earlier than did RIPK1 and Caspase-8 and disassociated from the RIPK3-complex at a later stage than did the other molecules (Fig. 4d). Similar results were observed for the MLKL-containing complex, where RIPK1 and Caspase-8 interacted with MLKL during PANoptosis, and this interaction was augmented by Z-VAD treatment. The interaction between RIPK3 and MLKL was detected after 30 min of treatment and was subsequently weakened (Fig. 4d). These results support the conclusion that the RIPK3-MLKL-RIPK1-Caspase-8 multimeric protein complex is formed during PANoptosis. Next, we immunoprecipitated Flag-RIPK3 from MLKL-depleted cell lysates after anti-V5 bead-mediated immunoprecipitation to obtain MLKL-free RIPK3-containing complexes. We observed the recruitment of RIPK1 and Caspase-8 to RIPK3, which did not bind to MLKL, indicating the presence of a RIPK3-RIPK1-Caspse-8 complex (Supplementary Fig. 4a). Given that RIPK3 forms a binary complex with MLKL, we performed immunofluorescence staining for RIPK3, RIPK1, and MLKL in the N-terminal V5-tagged MLKL-expressing PKLC-RIPK3^2×Fv^ cells treated with DZ to determine the presence of distinct RIPK3-containing protein complexes (Fig. 4e). Using immunofluorescence, we identified three RIPK3-containing protein complexes, namely RIPK3-MLKL, RIPK3-RIPK1, and RIPK3-MLKL-RIPK1, within a single cell (Fig. 4f, g). These findings prompted the investigation of whether these complexes are assembled independently and randomly. First, we determined the localization of RIPK1, RIPK3, MLKL, and Caspase-8, the primary components of the identified complexes, in TritonX-100-soluble and insoluble fractions. We observed significant changes in RIPK3 but not in MLKL, RIPK1, or pro-Caspase-8 in the detergent-resistant fraction, suggesting that MLKL, RIPK1, and pro-Caspase-8 could not form detergent-resistant amyloid-like complexes with RIPK3 in the TritonX-100-insoluble fraction (Supplementary Fig. 4b, c). In addition, RIPK3 did not translocate to the detergent-resistant fraction in the TSZ-treated cells, indicating that the RIPK3 aggregates were distinct from those in Dim- or DZ-treated cells (Supplementary Fig. 4d). To trace the dynamic state of RIPK3 in the DZ-induced complex, we restored N-terminally fused mRuby3-tagged RIPK3^2×Fv^ in WT MLKL- and N-terminal V5-tagged MLKL-expressing PKLCs. Live cell images showed similarly sized RIPK3 puncta in DZ-treated WT MLKL- and N-terminal V5-tagged MLKL-expressing cells (Supplementary Fig. 4e, f). The immunofluorescence of RIPK3 showed the same result, indicating that RIPK3 puncta increased over time (Supplementary Fig. 4g–i).

**Fig. 4.**
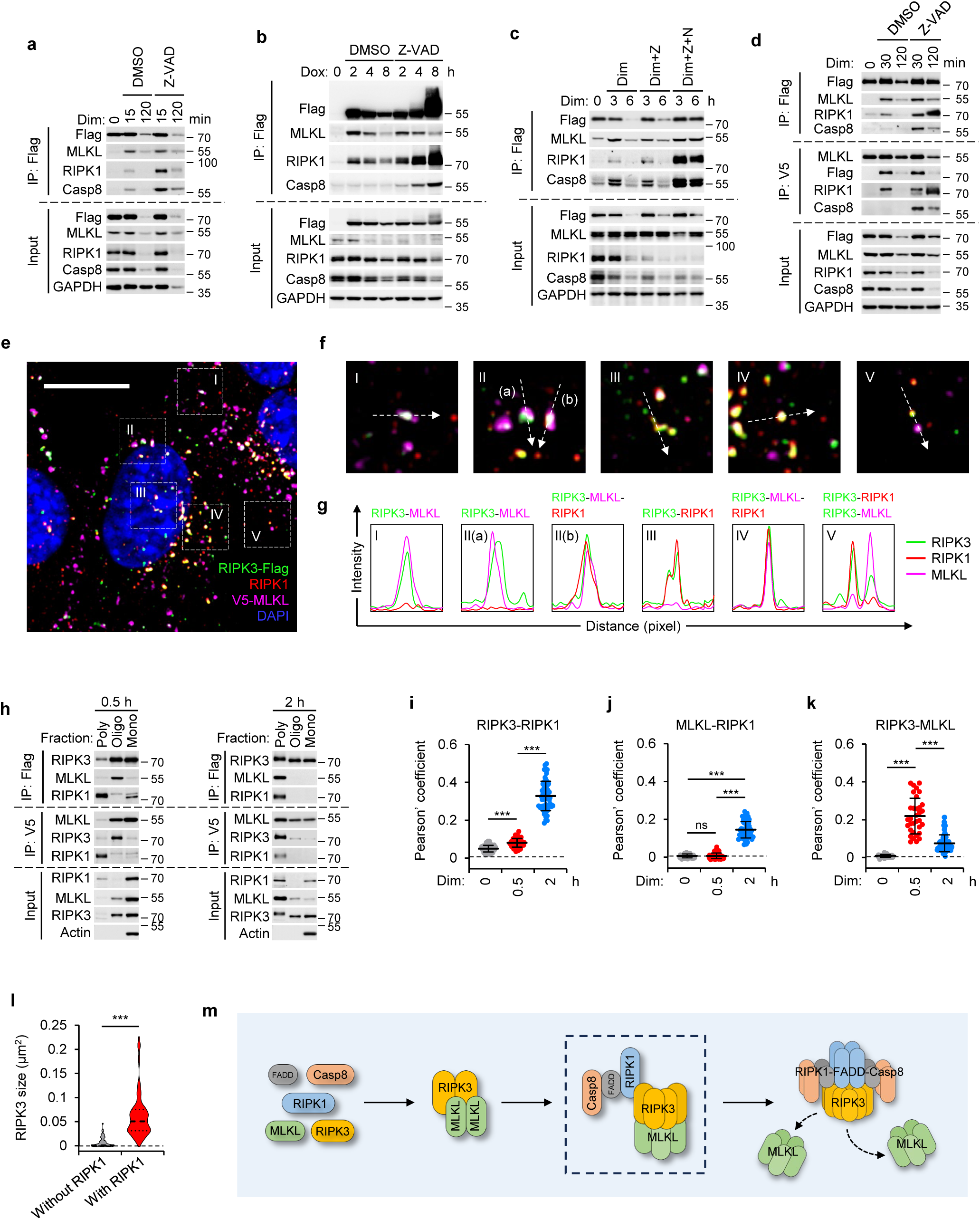
The affinity for RIPK3 coordinates stepwise assembly of MLKL and RIPK1 into the PANoptosome. **a-c** PKLC-RIPK3^2×Fv^ cells were treated with Dim + DMSO or Dim + Z-VAD (Z) for the indicated durations (**a**). PKLC-RIPK3^Tet-on^ cells were treated with Dox + DMSO or Dox + Z-VAD for the indicated durations (**b**). PL45-RIPK3^2×Fv^ cells were treated with Dim + DMSO, Dim + Z, or Dim + Z + NSA for the indicated durations (**c**). RIPK3 was immunoprecipitated from whole-cell lysates using anti-Flag resin. Lysates and immunocomplexes were analyzed by immunoblotting using antibodies against Flag, RIPK1, MLKL, and Casp8. **d** N-terminally V5-tagged MLKL-expressing PKLC-RIPK3^2×Fv^ cells were treated with Dim or Dim + Z-VAD for the indicated durations. RIPK3 or MLKL was immunoprecipitated from whole-cell lysates using anti-Flag or anti-V5 resin. Lysates and immunocomplexes were analyzed by immunoblotting using antibodies against Flag, RIPK1, MLKL, and Casp8. **e** Representative immunofluorescence image of PKLC-RIPK3^2×Fv^ cells stained with antibodies against Flag, V5-tag, and RIPK1 after treatment with Dim + Z for 2 hours. Scale bar, 10 μm. **f** Enlarged, cropped views of five selected areas from **e**. **g** Line graphs showing fluorescence intensity profiles of RIPK3, RIPK1, and MLKL along the representative dashed arrowheads in **f**. **h** N-terminally V5-tagged MLKL-expressing PKLC-RIPK3^2×Fv^ cells were treated with Dim + Z for 0.5 or 2 hours. Whole-cell lysates were fractionated using a Superose 6 Increase size-exclusion chromatography column. Fractions corresponding to monomers, oligomers, and polymers were collected and immunoprecipitated with anti-Flag or anti-V5 resin. Input fractions and immunocomplexes were analyzed by immunoblotting using the indicated antibodies. **i-k** Quantification of colocalization between RIPK3 and RIPK1 (**i**), RIPK1 and MLKL (**j**), or RIPK3 and MLKL (**k**) using Pearson correlation coefficients. **l** Quantification of the size of RIPK3 puncta with or without RIPK1 colocalization. **m** Schematic illustrating the RIPK3-PANoptosome assembly process. Data are presented as means ± SD. Statistical significance was determined by two-sided unpaired Student’s t-tests. ***p < 0.001; ns, no significance.

We collected the TritonX-100-soluble fraction from the DZ-treated cells and performed gel-filtration chromatography. As expected, RIPK3 and MLKL formed oligomers and polymers that were eluted in 13–16 and 9–10 mL fractions, respectively, 30 min and 2 h after treatment, revealing that the size of the RIPK3- or MLKL-containing complexes expanded in a time-dependent manner (Supplementary Fig. 4j). RIPK1 migrated directly to the polymer fractions 2 h after treatment but did not appear in the oligomer fractions (Supplementary Fig. 4j). To determine the distribution of the complexes in the different fractions, we collected 9–10 mL of eluent as the polymer fraction, 13–15 mL of eluent as the oligomer fraction, and 17–20 mL of eluent as the monomer fraction. Flag-RIPK3 and V5-MLKL were immunoprecipitated from each column fraction. Co-immunoprecipitation showed that MLKL bound to RIPK3 in the oligomer fraction at 30 min and in the polymer fraction at 2 h after treatment. In contrast to MLKL, RIPK1 interacted only with RIPK3 in the polymer fraction, regardless of the time point. Similar results were obtained using V5-MLKL immunoprecipitation (Fig. 4h). To confirm our observations, we performed pairwise co-immunostaining for RIPK3, MLKL, and RIPK1 and quantified the co-localization degree between these proteins. RIPK1 significantly co-localized with RIPK3 upon RIPK3 dimerization, and the co-localization markedly increased 2 h after treatment (Fig. 4i). The trend of RIPK1 and MLKL co-localization over time was consistent with that of RIPK1 and RIPK3 but was significantly weaker owing to the indirect interaction between RIPK1 and MLKL (Fig. 4j). Conversely, MLKL and RIPK3 showed a strong co-localization 30 min after treatment, which significantly decreased hours after treatment, consistent with MLKL disassociating from the RIPK3-complex at a later stage (Fig. 4k). This is consistent with the observation of numerous homopolymers of MLKL in previous triple immunofluorescence images (Fig. 4e). Additionally, larger RIPK3 puncta tended to recruit RIPK1 (Fig. 4l). We depicted the detailed process of a complex assembly during noncanonical PANoptosis, showing that the three complexes identified earlier represent different stages of a growing complex. RIPK3 dimerization triggered the gradual assembly of RIPK3 into polymers of increasing size (Fig. 4m). MLKL was rapidly recruited to low-polymerization RIPK3 oligomers and assembled into homopolymers on RIPK3 scaffolds to form the RIPK3-MLKL complex. With extending RIPK3 polymer, RIPK1 began to bind to RIPK3 to form the RIPK3-MLKL-RIPK1 complex. During RIPK3 growth and RIPK1 binding, mature MLKL homopolymers dissociated from the complex, leaving a RIPK3-RIPK1 binary complex. Current evidence suggests that ASC-dependent inflammasomes participate in the formation of canonical PANoptosomes, bridging necroptosis and apoptosis with pyroptosis^42^. Given that PKLCs lack the crucial molecules of the ASC-dependent inflammasome, we termed the RIPK3-MLKL-RIPK1-FADD-Caspase-8 complex dynamically assembled during RIPK3-PANoptosis as RIPK3-PANoptosome.

### RIPK3-RIPK1 structural configuration gates MLKL-mediated cytolytic execution

RIPK3-PANoptosome and TSZ-induced canonical necrosome have highly similar molecular compositions. Hence, we investigated the differences between the structures and functions of the two complexes. We monitored the phosphorylation kinetics of RIPK3, MLKL, and RIPK1 for up to 4 hours following DZ treatment or 8 hours post-TSZ exposure. DZ-induced phosphorylation of RIPK3 and MLKL occurred earlier than that induced by TSZ owing to direct RIPK3 activation. RIPK1 phosphorylation occurred earlier than those of RIPK3 and MLKL in TSZ-treated cells because RIPK1 activation is an upstream event during necrosome assembly. In contrast, RIPK1 phosphorylation was weaker and occurred later than those of RIPK3 and MLKL in DZ-treated cells (Fig. 5a), indicating that RIPK1 phosphorylation is a byproduct and dispensable for RIPK3-PANoptosis. These results were consistent with the previous observation that Nec-1s did not affect RIPK3-PANoptosis (Supplementary Fig. 3a, b). We subsequently compared the interaction between RIPK1 and RIPK3 in cells treated with DZ and TSZ by measuring the RIPK1 levels in RIPK3 immunoprecipitants. The binding between RIPK1 and RIPK3 in DZ-induced PANoptosomes was weaker than that in TSZ-induced necrosomes (Fig. 5b). To investigate the impact of this kinase-independent, weaker binding of RIPK1 on cell death, we monitored the changes in the ATP levels in PKLC-RIPK3^2×Fv^ cells over 6 h after DZ and TSZ treatments. DZ-induced necroptosis was attenuated at a later stage, with ATP levels recovering slowly, completely different from TSZ-induced necroptosis (Supplementary Fig. 5a, Fig. 5c). The phenomena of rapid decline in intracellular ATP and subsequent recovery were pronounced in RIPK1- or Caspase-8-deficient cells (Supplementary Fig. 5b, c). To investigate the differences in ATP dynamic changes between cells treated with DZ and TSZ, time-lapse phase contrast imaging was performed, which showed that Dim- and TSZ-treated cells underwent irreversible death within 24 h, while DZ-treated and Dim-treated RIPK1-deficient cells reattached and survived after rapidly rounding, revealing that the killing capacity of MLKL was severely weakened (Supplementary Fig. 5d, e). However, the phosphorylation levels of MLKL in DZ-treated cells were higher than those in TSZ-treated cells (Fig. 5a). These data prompted us to speculate that the MLKL-located molecular environment, and not the MLKL phosphorylation levels, affected its killing capacity. Next, we observed the structure of RIPK3 aggregates in DZ- and TSZ-treated PKLC-mRuby3-RIPK3^2×Fv^ cells using 3D structured illumination microscopy. High-resolution fluorescence images showed that RIPK3 formed small round puncta dispersed around the plasma and discontinuous along the z-axis in the DZ-treated cells (Fig. 5f). In TSZ-treated cells, RIPK3 puncta were prone to clustering and formed large amorphous fiber-like aggregates that were continuous along the z-axis (Fig. 5g). Previous studies have shown that, in TSZ-induced necrosomes, RIPK1 oligomerization initiates RIPK3 recruitment, forming heterogeneous amyloid fibrils and leading to necroptosis^43–45^. Our data revealed that, in contrast to the TSZ-induced complex, the DZ-induced PANoptosome began with RIPK3 oligomerization, which formed a small round complex with a low proportion of RIPK1 molecules bound to the surface, facilitating the exposure of the death domain by RIPK1 to recruit FADD and Caspase-8. Thus, the unique supramolecular architecture of the RIPK1-RIPK3 complex scaffolds enables co-activation of MLKL phosphorylation and caspase activation within individual cells during PANoptosis.

**Fig. 5.**
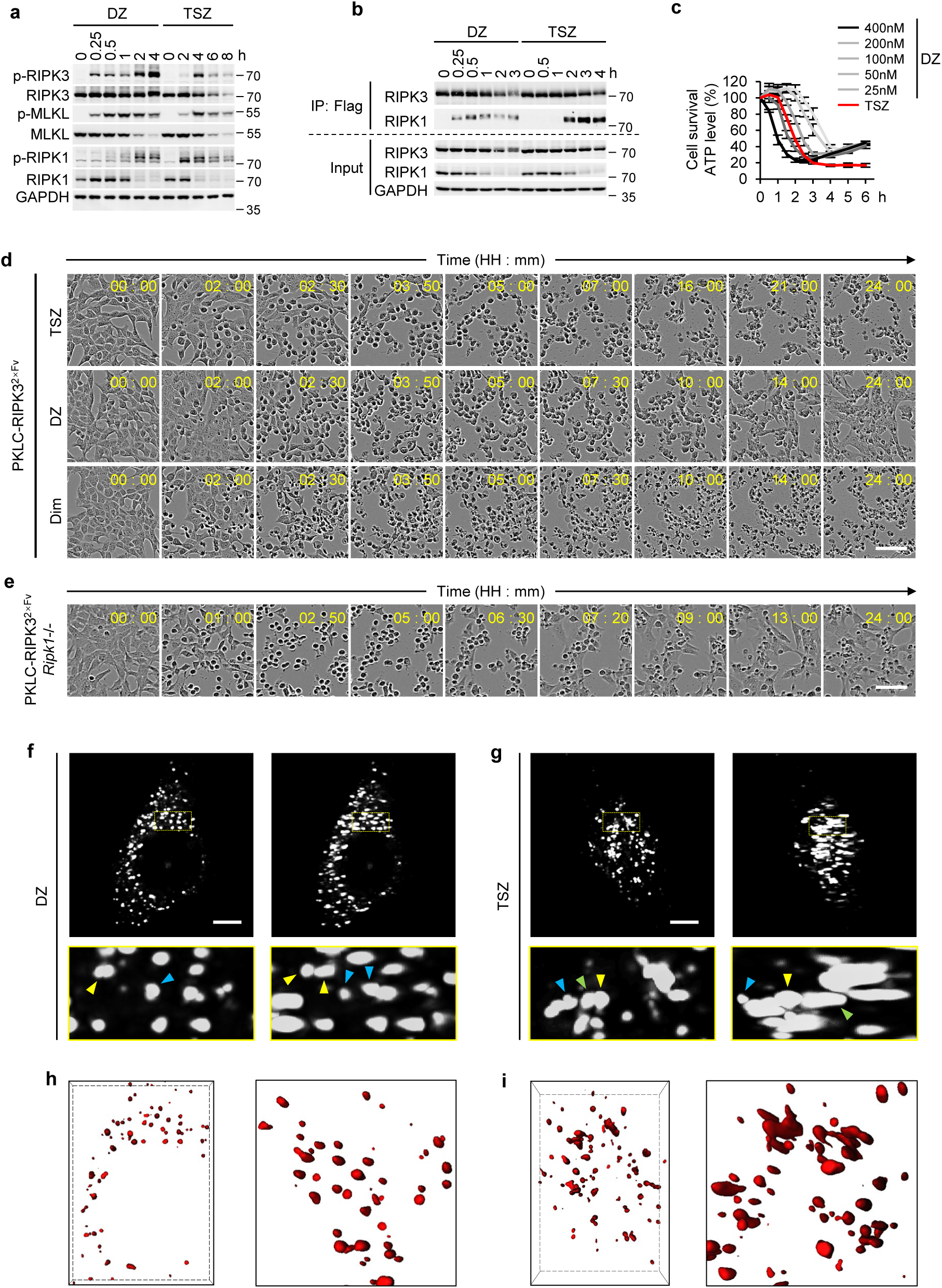
RIPK3–RIPK1 structural configuration gates MLKL-mediated cytolytic execution. **a, b** PKLC-RIPK3^2×Fv^ cells were treated with DZ (Dim, 50 nM) or TSZ (TNF, 40 ng/mL) for the indicated durations. RIPK3 was immunoprecipitated from whole-cell lysates using anti-Flag resin (**b**). Lysates and immunocomplexes were analyzed by immunoblotting using the specified antibodies. **c** PKLC-RIPK3^2×Fv^ cells were treated with TSZ or increasing concentrations of Dim + Z for the indicated durations. Cell viability was determined by measuring ATP levels using the CellTiter-Glo kit. **d, e** Phase-contrast time-lapse micrographs of PKLC-RIPK3^2×Fv^ (**d**) or RIPK1-deficient PKLC-RIPK3^2×Fv^ (**e**) cells treated with TSZ, DZ, or Dim alone. Images are representative of one of two fields from four independent experiments. Scale bar, 50 μm. **f, g** Representative live-cell 3D-SIM images of RIPK3-mRuby3 puncta in N-terminally V5-tagged MLKL-expressing PKLC-RIPK3^2×Fv^ cells treated with DZ for 2 hours (**f**) or TSZ for 4 hours (**g**). Rotated images around the Y-axis are displayed on the right. Arrows of the same color on the left and right indicate the same observed overlapping puncta located at different layers. Scale bar, 5 μm. **h, i** 3D surface models of DZ-induced (**h**) or TSZ-induced (**i**) RIPK3 puncta based on images in **f** and **g**.

### Caspase-mediated proteolytic processing of RIPK3 and GSDMD restricts cell lysis during RIPK3-PANoptosis

We previously concluded that caspase activation is negatively regulated by kinase-active RIPK3, a specific biomarker of necroptosis, revealing the impact of the necroptotic pathway on caspase processing. To investigate whether caspase activity also affects the necroptotic pathway, we compared the kinetics of ATP decline during RIPK3 dimerization-induced PANoptosis with or without a combination treatment with Z-VAD. The inhibition of caspase activity markedly enhanced PANoptotic cell death (Supplementary Fig. 5a). Additionally, Caspase-8- and RIPK1-deficient RIPK3^2×Fv^-PKLCs exhibited significantly increased cell deaths (Supplementary Fig. 5b, c). Combined with previous observations that the RIPK3^2×Fv^-Flag fusion protein was cleaved into a 40-kD Flag-tagged fragment simultaneously with Caspase-3 and PARP processing (Supplementary Fig. 2a, b, 3c, d, k, l, 4b, c), we presumed that the caspase-dependent cleavage of RIPK3 attenuates RIPK3-PANoptosis. We subsequently tested the biochemical markers of necroptosis, apoptosis, and pyroptosis via immunoblotting. Z-VAD treatment effectively blocked the cleavage of Caspase-3, GSDMD, and RIPK3 but significantly increased the relative phosphorylation levels of RIPK3 and MLKL, with upshifted migration bands of phospho-RIPK3 (Supplementary Fig. 5d). These observations were confirmed through immunoblotting using Caspase-8 knockout PKLC-RIPK3^2×Fv^ cells (Supplementary Fig. 5e). Furthermore, Z-VAD treatment did not affect cell death in Caspase-8-deficient RIPK3^2×Fv^-PKLCs (Supplementary Fig. 5f), indicating that Z-VAD functions through catalytic inhibition.

RIPK3 could be cleaved by caspase at Asp333^46–49^. To explore the functional significance of this cleavage event, we introduced a caspase-resistant point mutation at Asp333. We generated PKLC-RIPK3-D333E^2×Fv^ cell lines expressing the caspase-resistant RIPK3-D333E^2×Fv^ protein. Cell death induced by the dimerization of the RIPK3-D333E mutant was faster than that induced by WT RIPK3 (Supplementary Fig. 5g). Immunoblotting revealed increased RIPK3 and MLKL phosphorylation (Supplementary Fig. 5h). Although the RIPK3-D333E mutant could not be cleaved, its dimerization-induced caspase activation and apoptosis in MLKL-deficient PKLCs remained at the same levels as those in WT RIPK3-expressing cells (Supplementary Fig. 5i, j). This is consistent with previous finding that expression of RIPK3^D333∼end^ truncation could directly induce caspase activation (Supplementary Fig. 3m).

The cleavage of RIPK3 and RIPK1 during PANoptosis was mediated by Caspase-3 and was completely blocked by Caspase-3 deficiency (Supplementary Fig. 5k). Furthermore, treatment with Caspase-3-specific inhibitor Z-DEVD-FMK effectively inhibited RIPK3 cleavage at concentrations that did not affect apoptosis in Dim-treated MLKL/GSDMD-double deficient PKLCs (Supplementary Fig. 5l, m), suggesting that RIPK3 is only cleaved by highly active Caspase-3.

As the Asp to Ala mutation is common in cleavage-resistance studies, we first measured necroptotic cell death induced by the DZ and TSZ after introducing D333A mutation in PKLC-RIPK3^2×Fv^ cells. The cell death of all RIPK3-D333A^2×Fv^-expressing PKLC clones was alleviated (Supplementary Fig. 6a, b). Similarly, we observed strongly reduced RIPK3 and MLKL phosphorylation (Supplementary Fig. 6c, d), identifying RIPK3-D333A as a mutant with weakened kinase activity. In the MLKL-deficient PKLC-RIPK3-D333A^2×Fv^ clones, Dim-induced apoptotic cell death was significantly enhanced (Supplementary Fig. 6e). The cleavage of Caspase-8, Caspase-3, and downstream GSDMD was greatly enhanced in MLKL-expressing and -deficient PKLC-RIPK3-D333A^2×Fv^ cells compared with that in WT PKLC-RIPK3^2×Fv^ cells (Supplementary Fig. 6d, f), indicating that RIPK3 kinase activity negatively regulates caspase activation.

In contrast to Caspase-8 deficiency, Caspase-3 knockdown did not block caspase-dependent cell death in MLKL-deficient PKLCs (Supplementary Fig. 7a). Therefore, we traced the ATP decline and intracellular lactate dehydrogenase (LDH) release in MLKL-deficient and MLKL-/Caspase-3-double deficient cells over 7 h after Dim treatment. Caspase-3 knockdown had a slight effect on ATP decline (Supplementary Fig. 7b) but strongly increased the LDH release caused by caspase-dependent cell death (Supplementary Fig. 7c), suggesting that Caspase-3 knockout could switch apoptosis to cell lytic death. Our previous investigation showed weak and unstable GSDMD p30 but strong p20 fragment signals upon caspase activation, revealing negative cleavage of the GSDMD active p30 fragment. Growing evidence suggests that Caspase-3 cleaves GSDMD within its N-terminal domain, yielding p20 fragments that do not mediate pyroptosis^50,51^. Our immunoblotting data confirmed that Caspase-3 knockdown blocked secondary processing (Supplementary Fig. 7d) and enhanced the membrane accumulation of the GSDMD p30 fragment (Supplementary Fig. 7e). GSDMD knockdown entirely blocked caspase-dependent cell death in MLKL-/Caspase-3-double deficient cells (Supplementary Fig. 7f). In addition, Dim-induced cell death in MLKL-/Caspase-3-double deficient cells displayed classical pyroptotic features, including cell rounding, bubble-like swelling, and membrane rupture. The nuclei of pyroptotic cells collapsed, causing propidium iodide (PI)-positive chromatin to disperse into bubble-like cell swellings (Supplementary Fig. 7g). Similarly, GSDMD-D88A, a Caspase-3-resistant mutant, mediated enhanced LDH release but not ATP decline in Caspase-3-sufficient cells upon RIPK3 dimerization (Supplementary Fig. 7h–k). As RIPK3-PANoptosis is independent of ASC and Caspase-1, we hypothesized that the primary processing of GSDMD is mediated by Caspase-8, as recent discoveries have suggested that Caspase-8 directly cleaves GSDMD to trigger pyroptosis^51–56^. Consistent with this, Caspase-8 knockdown completely blocked caspase-dependent cell death (Supplementary Fig. 7l) and GSDMD cleavage in MLKL-/Caspase-3-double deficient cells (Supplementary Fig. 7m).

### RIPK1 kinase-dependent priming licenses RIPK3 activation followed by kinase-independent recruitment executing caspase engagement

Given the caspase-dependent cleavage of RIPK1 observed during RIPK3-PANoptosis and its dispensable role in MLKL phosphorylation, we examined whether RIPK1 cleavage regulates apoptosis in MLKL-deficient cells. First, we evaluated cell death induced by lentivirus-mediated overexpression of WT RIPK1 and RIPK1-D325A, a caspase-resistant mutant, in MLKL-/RIPK1-double deficient cells (Supplementary Fig. 8a). RIPK1-D325A caused approximately 46% cell death 36 h after viral infection, whereas approximately 100% of the WT RIPK1-expressing cells survived. RIPK1-D325A-expressing cells died faster than did WT RIPK1-expressing cells 60 h after viral infection, and this was blocked by Z-VAD treatment (Supplementary Fig. 8b). Additionally, Caspase-3 and −8 cleavage was significantly enhanced by introducing the D325A mutation (Supplementary Fig. 8c), indicating that caspase-dependent cleavage of RIPK1 inhibits the activation of the apoptotic pathway. The overexpression of WT RIPK1 and RIPK1-D325A failed to induce cell death in RIPK3-null PKLCs (Supplementary Fig. 8d). Thus, RIPK1 overexpression possibly recruits RIPK3, facilitating RIPK3 polymerization, which recruits RIPK1 to form the RIPK1-RIPK3-RIPK1 complex, mediating caspase-dependent cell death. Since RIPK1-D325A-expressing PKLCs were sensitive to caspase-dependent cell death (Supplementary Fig. 8b, d), we failed to generate a cell line that constitutively expressed RIPK1-D325A.

We previously observed that the RIPK1^D325∼end^ failed to induce cell death without RIPK3 dimerization (Supplementary Fig. 3e, 8d), revealing the significance of the RIPK1 kinase domain in mediating RIPK1-initiated cell death. However, Nec-1s completely blocked RIPK1-induced cell death in RIPK3-expressing PKLCs (Supplementary Fig. 8e), confirming the importance of RIPK1 kinase activity in RIPK1-initiated PANoptosis. Kinase-active RIPK1 recruited RIPK3 as a seed, mediating RIPK3 priming and triggering RIPK3-PANoptosis (Supplementary Fig. 8f).

### RIPK3–PANoptosis generates unique cytomorphological signatures during cell death execution

PANoptosis is poorly understood owing to the lack of detailed morphological characterization. PKLC-RIPK3^2×Fv^ cells underwent necroptosis and apoptosis after treatment with DZ and DG, respectively. Pyroptosis was also induced by Dim treatment in MLKL- and Caspase-3-deficient cell lines. This allowed us to induce four PCD types with the same stimuli in a single parental cell line, enabling an accurate comparison of the morphological differences among these PCDs. As expected, DG-treated cells showed typical apoptotic morphology, such as membrane blebbing, cell shrinkage, and fragmentation into membrane-bound apoptotic bodies (Fig. 6a). DZ-induced necroptosis began with cell rounding and swelling, followed by the loss of cell content and PI uptake, while the nucleus remained intact (Fig. 6a). Dim-induced pyroptosis in MLKL- and Caspase-3-deficient cells resulted in cell rounding, bubble-like swelling, and membrane rupture. In contrast to the nuclear condensation observed in apoptosis, the nuclei of pyroptotic cells collapsed, causing PI-positive chromatin to disperse into bubble-like cell swelling (Supplementary Fig. 7g). PANoptosis began with sequential cell rounding and blebbing, combining the early morphological features of apoptosis, necroptosis, and pyroptosis. Following cell blebbing, the cytomatrix divided into numerous small membrane-bound vesicles that accumulated on one side of the cell (Fig. 6b). The nuclei and organelles were tightly compressed on the other side of the cell, resulting in the formation of PANoptotic cells with a nomad jellyfish-like structure (Fig. 6b, c). Subsequently, a large bubble emerged from the cell surface and connected to many small vesicles derived from the cytoplasmic matrix. The plasma membrane integrity was lost, accompanied by the disintegration of the nuclear membrane, causing the release of cell contents and the dispersion of PI-stained chromatin into cell bubbles and small vesicles on the cell surface (Fig. 6b). Notably, organelles, such as mitochondria, were confined to a certain area within dead cells (Fig. 6c). Based on these morphological changes, PANoptosis was divided into five distinct stages (Fig. 6b, d). These observations revealed significant morphological differences between PANoptosis and the other PCD types.

**Fig. 6.**
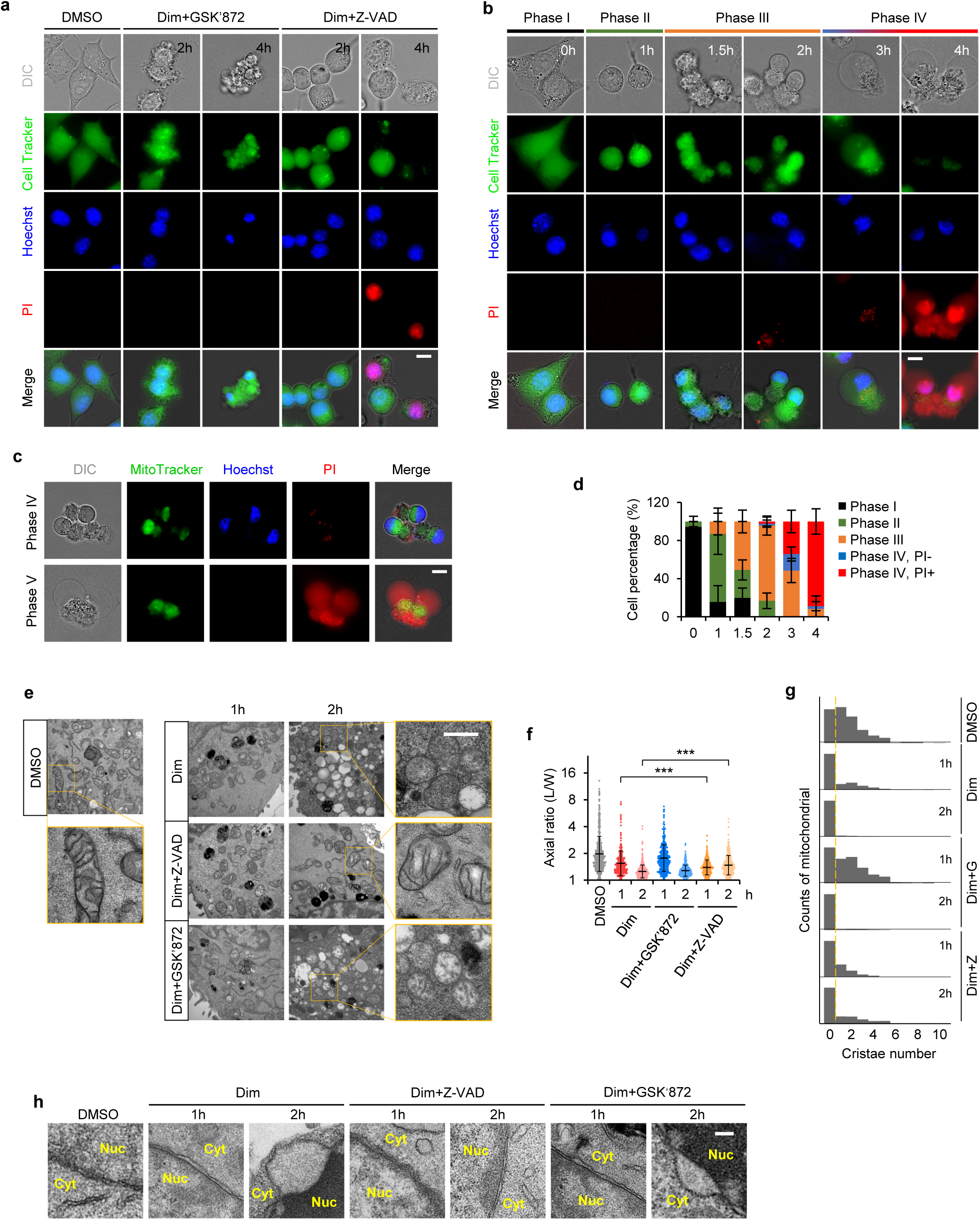
RIPK3–PANoptosis generates unique cytomorphological signatures during cell death execution. **a, b** Representative images of PKLC-RIPK3^2×Fv^ cells treated with Dim + GSK’872, Dim + Z-VAD (**a**) or Dim (**b**) for the indicated durations. Cell morphology was visualized by wide-field light microscopy, and plasma membrane integrity was monitored by propidium iodide (PI) uptake and CellTracker release. Nuclei were labeled with Hoechst. PANoptotic cell death was categorized into five stages (Phase I–IV) based on morphological features at different timepoints (**b**). Scale bar, 10 μm. **c** Representative images of mitochondrial localization in Phase IV–V PANoptotic cells. Mitochondria and nuclei were labeled with MitoTracker Green and Hoechst, respectively. Plasma membrane integrity was monitored by PI uptake and CellTracker release. **d** Quantification of the percentage of cells in Phase I–IV from ten fields at each timepoint in **b**. Data are presented as means ± SD. **e-g** Representative TEM images of mitochondria in PKLC-RIPK3^2×Fv^ cells treated with Dim, Dim + Z-VAD, or Dim + GSK’872 for 1 or 2 hours. Axial ratios (**f**) and cristae number distributions of mitochondria (**g**) from ten randomly selected cells were quantified. Statistical significance was determined by two-sided unpaired Student’s t-tests. Data are presented as means ± SEM. ***p < 0.001(**g**). **h** Representative TEM images of nuclear envelopes in PKLC-RIPK3^2×Fv^ cells treated with Dim, Dim + Z-VAD, or Dim + GSK’872 for 1 or 2 hours.

The mitochondria are central organelles in metabolism and energy conversion and multifaceted regulators of PCDs. The morphological structures of the mitochondria are regulated by PCDs, which affects the process of cell death and its outcomes^57^. Hence, we observed morphological changes in the mitochondria during RIPK3-PANoptosis using scanning electron microscopy. After Dim treatment, the mitochondria gradually became rounded from an elongated morphology (Fig. 6e), with a decreased axial ratio (Fig. 6f), accompanied by the loss of cristae and matrix vacuolation (Fig.6 e, g). The mitochondria of the cells under DG-induced apoptosis underwent a similar process but more slowly (Fig. 6e–g). These results suggest that the mitochondrial damage was primarily mediated by phospho-MLKL in the early stages of PANoptosis, which was further confirmed by the rapid rounding of mitochondria during DZ-induced necroptosis. Similarly, the tendency for further rounding of the mitochondria and crest loss was blocked or even reversed by adding Z-VAD (Fig. 6 e–g), suggesting that mitochondrial damage is primarily mediated by caspases in the later stage of PANoptosis, consistent with the sequential activation of MLKL and caspases. The dissociation of the nuclear envelope and the blebbing of the outer membrane were observed in the PANoptotic cells; however, these were blocked by GSK’872 but not Z-VAD (Fig. 6h), suggesting that the changes in the nuclear envelope are mediated by caspases. This may be related to the disintegration of the nucleus, as previous data revealed that nuclear disruption occurred in Dim-induced PANoptosis or DG-induced apoptosis and in Dim-induced pyroptosis of MLKL-/Caspase-3-double deficient PKLC-RIPK3^2×Fv^ cells, where PI-positive nuclear DNA diffused into the cytoplasm or apoptotic bodies, while the nuclear envelope remained integrated in DZ-induced necroptosis (Fig. 6a, b, Supplementary Fig. 7g).

### RIPK1 dominantly coordinates chemokine production and macrophage recruitment via NF-κB during PANoptosis execution

Necroptosis drives cytokine production in a MLKL-^58–60^ or RIPK1-kinase-^61^dependent manner. Canonical PANoptosis and pyroptosis drive IL-1β secretion through Caspase-1-mediated pro-IL-1β processing^42^. To characterize cytokine secretion in RIPK3-PANoptotic cells, we profiled the transcriptome of Dim-treated PKLC-RIPK3^2×Fv^ cells via RNA sequencing. Based on a differential gene expression analysis, we identified a transcriptional signature of 43 cytokines produced during RIPK3-PANoptosis and found that the most highly upregulated cytokines belonged to chemokines from the CXCL*-* and CCL*-* families (Fig. 7a). We further analyzed the protein levels of cytokines released from PANoptotic cells using Luminex assay. The induction of RIPK3-PANoptosis led to increased secretion of four chemokines, namely CXCL1, CXCL10, CCL2, and CCL20, into the conditioned media (Fig. 7b). These chemokines were also upregulated at the transcriptional level, which was confirmed via real-time RT-PCR in both mouse PKLC-RIPK3^2×Fv^ and human PL45-RIPK3^2×Fv^ cells (Fig. 7c, d). Notably, high levels of CXCL5 were released into the media in response to PANoptosis induction, whereas the mRNA levels of *Cxcl5* showed slight upregulation (Fig. 7a, b), indicating that CXCL5 release was not regulated by transcription efficiency.

**Fig. 7.**
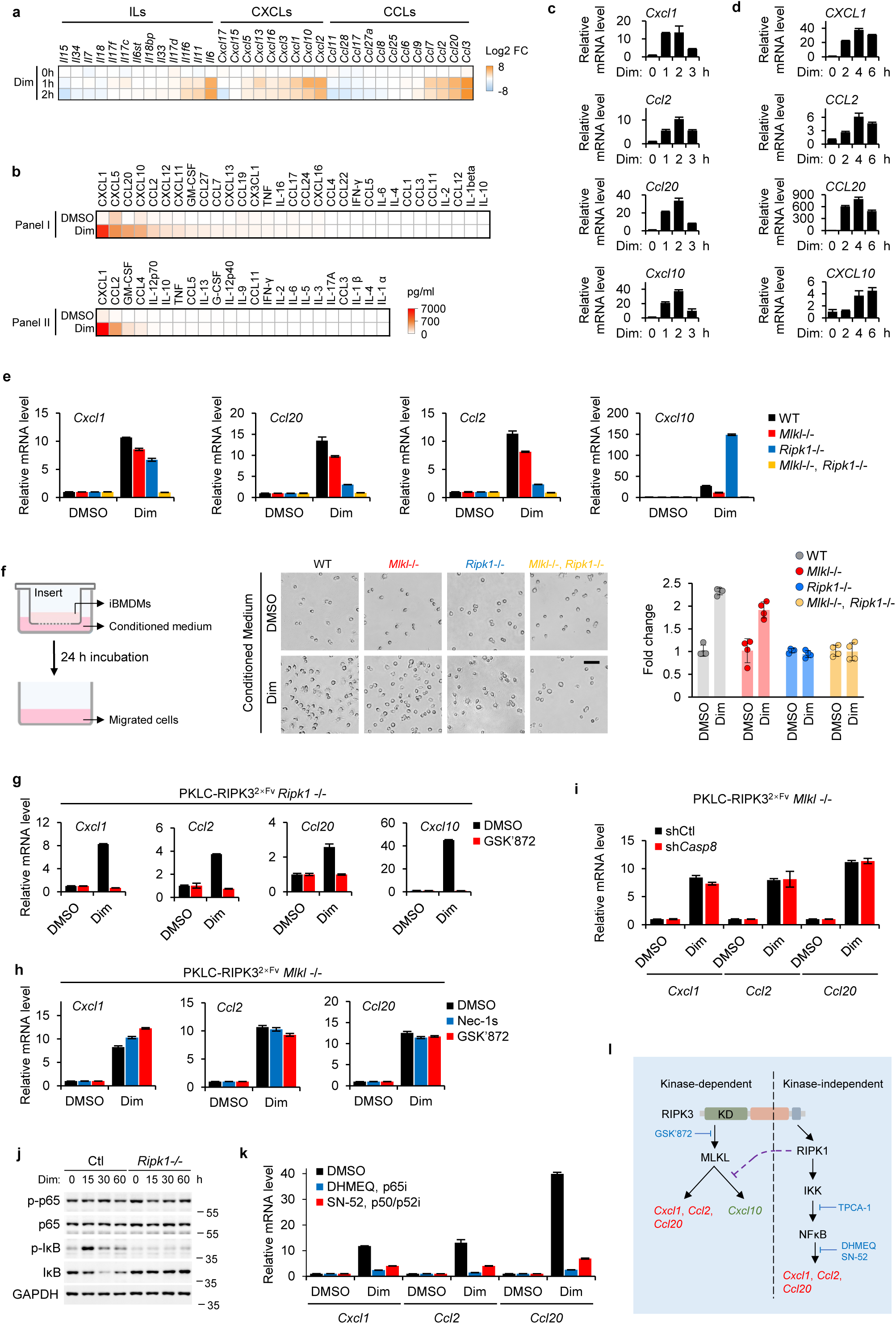
RIPK1 dominantly coordinates chemokine production and macrophage recruitment via NF-κB during PANoptosis execution. **a** Expression profiles (log2-based fold change of FPKM) for cytokines in PKLC-RIPK3^2×Fv^ cells treated with Dim for the indicated durations. **b** Secreted cytokine levels in conditioned media from Dim-treated PKLC-RIPK3^2×Fv^ cells, measured by Luminex assay. **c, d** PKLC-RIPK3^2×Fv^ (**c**) and PL45-RIPK3^2×Fv^ (**d**) cells were treated with Dim for the indicated durations. Relative mRNA levels of *CXCL1*, *CCL2*, *CCL20*, and *CXCL10* were quantified by qRT-PCR. **e** qRT-PCR quantification of *CXCL1*, *CCL2*, *CCL20*, and *CXCL10* mRNA levels in PKLC-RIPK3^2×Fv^ cells (with indicated genotypes) treated with 50 nM Dim for 1 hour. **f** Immortalized bone marrow-derived macrophages (iBMDMs) migrated toward conditioned medium (CM) from living or Dim-treated PKLC-RIPK3^2×Fv^ cells (with indicated genotypes). The fetal bovine serum (FBS) in all CM was pre-inactivated before PKLCs culturing. iBMDMs were added to the upper insert, and migrated cells in the bottom well were monitored by microscopy. Scale bar, 50 μm. **g** qRT-PCR quantification of *Cxcl1*, *Ccl2*, and *Ccl20* mRNA levels in RIPK1-deficient PKLC-RIPK3^2×Fv^ cells treated with 50 nM Dim ± GSK’872 for 1 hour. **h** qRT-PCR quantification of *Cxcl1*, *Ccl2*, *Ccl20*, and *CXCL10* mRNA levels in MLKL-deficient PKLC-RIPK3^2×Fv^ cells treated with 50 nM Dim ± GSK’872 or ± Nec-1s for 1 hour. **i** qRT-PCR quantification of *Cxcl1*, *Ccl2*, and *Ccl20* mRNA levels in WT and caspase-8 knockdown PKLC-RIPK3^2×Fv^ cells treated with 50 nM Dim for 1 hour. **j** Immunoblotting analysis of NF-κB activation markers in MLKL-deficient PKLC-RIPK3^2×Fv^ cells (WT or RIPK1-deficient) treated with Dim for the indicated durations. Results are representative of three independent experiments. **k** qRT-PCR quantification of *Cxcl1*, *Ccl2*, and *Ccl20* mRNA levels in MLKL-deficient PKLC-RIPK3^2×Fv^ cells treated with 50 nM Dim ± DHMEQ or ± SN-52 for 1 hour. **l** Hypothesis illustrating parallel pathways regulating chemokine production during RIPK3-PANoptosis via kinase-dependent or -independent mechanisms. Data are presented as means ± SD of triplicate wells (**c-e, g-i, k**).

Subsequently, we examined the involvement of the PANoptotic machinery in the production of these chemokines. We found that MLKL and RIPK1 knockout reduced the production of CXCL1, CCL2, and CCL20. However, the effects of RIPK1 depletion were more robust. In addition, the double knockout of MLKL and RIPK1 completely blocked the upregulation of CXCL1, CCL2, and CCL20 (Fig. 7e). Thus, although CXCL1, CCL2, and CCL20 production relies on two parallel axes that depend on MLKL or RIPK1, RIPK1 plays a major role. In contrast, RIPK1 ablation strongly inhibited the transcription of CXCL10, which depends on MLKL (Fig. 7e), indicating that the MLKL-induced production of CXCL10 has a different regulatory mechanism from that of CXCL1, CCL2, and CCL20. Macrophages are attracted to cytokine secreted during cell death. Hence, we used conditioned media (CM) collected from WT, MLKL-, RIPK1-, or MLKL/RIPK1-knockout PKLC-RIPK3^2×Fv^ cultures to test their potency to mediate macrophage migration using a trans-well assay. Consistent with chemokine production, the CM from dying MLKL-knockout cells exhibited a slightly low ability to attract macrophages; however, following RIPK1 depletion, the CM from pure necroptosis failed to promote macrophage migration (Fig. 7f). These results demonstrate the crucial role of RIPK1 in promoting macrophage migration to dying cells.

In addition, the inhibition of RIPK3 kinase activity blocked the MLKL-dependent production of CXCL1, CXCL10, CCL2, and CCL20 in RIPK1-knockout PKLC-RIPK3^2×Fv^ cells (Fig. 7g), as previously reported^58,59^. RIPK1-dependent chemokine induction in MLKL-knockout cells was not inhibited by RIPK1 or RIPK3 kinase inhibitor (Fig. 7h). Since RIPK1 induces caspase-associated cell death in a kinase-independent manner, we investigated whether RIPK1-dependent chemokine induction relies on its cell death function. To address this, we examined CXCL1, CCL2, and CCL20 production in Caspase-8-depleted MLKL-null PKLC-RIPK3^2×Fv^ cells. Caspase-8 knockdown using shRNA did not abrogate RIPK1-dependent chemokine expression following the induction of PANoptosis (Fig. 7i).

KEGG pathway analysis of the Dim- vs. DMSO-treated PKLC-RIPK3^2×Fv^ cells revealed the enrichment for differentially expressed genes with annotations for NF-κB, MAPK, and PI3K-Akt signaling pathways (Supplementary Fig. 9a). Subsequently, we confirmed whether the identified pathways were activated during RIPK3-PANoptosis. RIPK3 activation led to the increased phosphorylation of JNK, p38, IKK, PI3K, and AKT, which was inhibited by RIPK1 ablation (Supplementary Fig. 9b). However, MEK and ERK phosphorylation declined rapidly after PANoptosis induction, and the altered ERK phosphorylation was not regulated by RIPK1 (Supplementary Fig. 9b). To examine the involvement of these pathways in RIPK1-mediated chemokine production, we evaluated the potential roles of IKK, PI3K, AKT, and MAPKs, such as JNK, MEK and p38, by screening several inhibitors. JNK (SP600152, JNK-in-8 and Tanzisertib), MEK (RO4987655 and Refametinib), ERK (SCH772984), PI3K (LY294002 and Alpelisib), and AKT (Capivasertib and AKT inhibitor VIII) inhibitors did not block CXCL1, CCL2, or CCL20 production during PANoptosis (Supplementary Fig. 9c–f). The addition of the IKK inhibitor TPCA-1 abolished the induction of chemokines, suggesting that IKK activation is required for RIPK1-dependent chemokine expression during RIPK3-PANoptosis (Supplementary Fig. 9g). Moreover, the inhibition of p38 activity by Ralimetinib or Losmapimod caused a partial reduction in chemokine expression (Supplementary Fig. 9g), indicating a regulatory role of p38 in mediating RIPK1-dependent chemokine induction. Thus, we focused on NF-κB activation. Dim stimulation led to the phosphorylation of p65 and IκB, as well as IκB degradation, vital signal transduction events for NF-κB activation (Fig. 7j). Notably, these events were inhibited by RIPK1 knockout (Fig. 7j). Subsequently, we blocked the nuclear translocation of p65 by DHMEQ or p50 by SN-52, critical events for NF-κB activation. Similar to the inhibition of the upstream kinase IKK, the inhibition of the nuclear translocation of NF-κB abolished chemokine induction (Fig. 7k).

These results suggest that chemokine production during PANoptosis can be induced in RIPK kinase-dependent and -independent manners. The kinase-dependent expression of chemokines relies on MLKL phosphorylation, which plays a minor role. However, the kinase-independent expression of chemokines relies on the non-cell death function of RIPK1, requiring IKK-mediated activation of NF-κB, which dominantly regulates chemokine production and macrophage recruitment (Fig. 7l).

## Discussion

The canonical activation of RIPK3 depends on RHIM-RHIM interactions between RIPK3 and various upstream adaptors. However, the pathologically increased expression of RIPK3^62^, the depletion of the negative regulator of RIPK3 amyloid aggregates^63^, or NHE1-mediated cytosol alkalization^37^ activate RIPK3 without other RHIM adaptors in a noncanonical manner, inducing MLKL-dependent necroptosis. Here, we found that, in several cell lines, the direct activation of RIPK3 induced MLKL phosphorylation and caspase cleavage, and activated caspases further cleaved gasdermin family members if expressed. For the first time, we observed that MLKL phosphorylation and caspase cleavage occur within the same cell upon OS-induced RIPK3 activation, chemical-induced dimerization of RIPK3, or artificial ectopic overexpression of RIPK3, which is direct evidence of PANoptotic cell death.

Although sensors upstream of PANoptosis have been discovered recently, the assembly mechanism of PANoptosomes remains poorly understood^30–35^. Because canonical PANoptosis relies on ASC-speck, with a complicated composition and architecture, investigating the molecular mechanism of PANoptosome assembly is challenging^42^. In this study, we identified a novel RIPK3-PANoptosome without the ASC complex, which has a relatively simple composition. Thus, using an artificial RIPK3-dimerization system, we constructed a sequential recruitment model for PANoptosome assembly. In this model, MLKL and RIPK1 recruitments were not mutually exclusive. They bound to RIPK3 sequentially to form different complexes, including the RIPK3-MLKL-RIPK1 ternary complex in the intermediate state, suggesting that all these dynamically assembled complexes belong to RIPK3-PANoptosomes. We also revealed, for the first time, the difference in the affinities of MLKL and RIPK1 for RIPK3, which led to the sequential recruitment and activation of MLKL and RIPK1 during RIPK3-PANoptosis.

Another significant finding of this study is that the killing activity of MLKL is not solely determined by its phosphorylation level; it is also influenced by the complex environment of MLKL. Death receptor-mediated activation of RIPK3 usually results in the formation of RIPK1-RIPK3 heterooligomers, in which RIPK1 and RIPK3 chimerize to form a large fibrous precipitate^43–45^, potentiating the strong killing activity of MLKL. However, after direct activation, RIPK3 forms scattered punctate homo-aggregates that phosphorylate MLKL more intensely, while MLKL toxicity remains low. Under this condition, cells rapidly become round, and intracellular ATP levels decline, entering a sublethal state but ultimately surviving. This suggests that the ATP decline during MLKL-mediated cell death is not only attributed to the ATP release resulting from plasma membrane disruption and the extracellular ATPase-catalyzed hydrolysis. Phospho-MLKL causes rapid depletion of intracellular ATP in the early stages of necroptosis, preceding plasma membrane permeabilization; however, this requires further investigation.

We propose that PANoptosis is a homeostatic system regulated by different pathways that collectively determine the process and consequences of cell death. The three activated PCD pathways involved in PANoptosis are interconnected and crucial. MLKL- and GSDMD-mediated membrane rupture is negatively regulated by the Caspase-3-dependent cleavage of RIPK3 and GSDMD-p30 fragments. Genetic ablation of apoptosis-associated molecules, such as Caspase-8 and RIPK1, or chemical inhibition of caspase activity enhanced MLKL- and GSDMD-mediated cell lysis. RIPK3 kinase activity is vital in triggering necroptosis; however, it markedly hinders RIPK1 recruitment and alleviates caspase activation.

In this study, for the first time, we observed the morphological features of PANoptosis extensively, which differ from those of apoptosis, pyroptosis, and necroptosis. Our limitation lies in not observing this specific morphology in other cells, potentially because of the varying proportions of MLKL and RIPK1 in different cells. In PKLCs, by the ectopic expression of RIPK1 or MLKL, we discovered that when the MLKL levels were increased, cells tended to exhibit classical necroptosis morphology, whereas increased RIPK1 levels favored apoptosis (data not shown). However, in these cell lines with different RIPK1/MLKL ratio, multiple cell death pathways were simultaneously activated and influenced each other. Therefore, relying on morphological features to identify the type of cell death is imprecise. Our morphological observations illustrate a potential cell morphology where necrosis and apoptosis exist in a delicate balance, enhancing the understanding of cell death morphology.

Necroptosis frequently triggers an inflammatory response through the transcriptional upregulation of various cytokines, a process dependent on MLKL^58–60^. Additionally, RIPK1 activates the JAK-STAT1 pathway via its kinase activity, mediating chemokine expression^61^. During RIPK3-induced PANoptosis, four chemokines (CXCL1, CCL2, CCL20, and CXCL10) were significantly upregulated. Notably, CXCL10 upregulation relied on phosphorylated MLKL, which was negatively regulated by RIPK1. In contrast to that of CXCL10, the upregulation of the other three chemokines relied on MLKL and RIPK1 independently, with the RIPK1-dependent pathway predominating, potentially owing to the restricted activity of MLKL within the PANoptosome. These RIPK1-dependent pathways operate independently of its kinase activity or the downstream activation of caspases, a mechanism that has not been reported. Inhibitor screening revealed that the RIPK1 kinase-independent upregulation of chemokines was driven by IKK-mediated NF-κB activation. However, the mechanism by which IKK is activated downstream of RIPK1 remains unclear. Our data underscore the critical role of RIPK1 in the inflammatory response, which contributes to the stronger immunogenicity of PANoptosis than that of necroptosis.

Our results indicate that the direct activation of RIPK3-mediated PANoptosis depends on the cell type. In many cell types, the direct activation of RIPK3 predominantly induces necroptosis without detectable caspase cleavage. This suggests that unknown regulatory mechanisms underly the assembly and outcome of RIPK3-initiated PANoptosis. Furthermore, we used a few cancer cell lines to simplify the PANoptosis model, enabling a focused investigation of the core molecular mechanisms. However, identifying PANoptosis-associated disease models remains an essential focus for future research.

## Methods

### Cell culture

786-O, PL-45, LLC, NIH-3T3, iBMDM, HEK293T and PKLC cells were cultured in Dulbecco’s modified Eagle’s medium (DMEM) supplemented with 10% fetal bovine serum (FBS) and 100 units/mL penicillin/streptomycin. All cells were cultured at 37 °C in a 5% CO_2_ incubator. All cell lines were tested to be mycoplasma-negative by the standard RT-PCR method.

### Cell death assay

1.8×10^4^ PKLCs were seeded into each well of 96-well plate and allowed to grow for 12 h, and then treated with the indicated compounds for different periods of time. To monitor the survival curve of PKLCs, cells were treated with B/B homodimerizer, Dox or TSZ every 30-, 45- or 60-min. The intracellular ATP levels of the remaining cells were measured to determine cell viability with the CellTiter-Glo Luminescent Assays kit (Promega) according to the manufactory’s instructions. In brief, 25 μL of CellTiter-Glo reagent was added to the cell culture medium and the cells were incubated with shaking at room temperature for 15 min. Luminescent recording was performed with Synerg H1 plate reader (BioTek).

### Morphological characterization of cell death

PKLCs were seeded in 29-mm glass-bottom dishes coated with 0.1 mg/mL poly-D-lysine and allowed to grow to about 80% confluence, and then treated with DMSO or 100 nM of B/B homodimerizer in combination with or without indicated inhibitors. Plasma membrane integrity was monitored by propidium iodide (PI) uptake and CellTracker or EGFP release. Nucleus and mitochondria were labeled by Hoechst 33342 (Invitrogen) and MitoTracker Green respectively. The morphological changes and plasma membrane permeability were observed and imaged using a BZ-X810 fluorescence microscope (KEYENCE). For Phase-contrast time-lapse imaging, PKLCs were seeded in 96-well plated as indicated. Cells were treated with 100 nM Dim or DZ for 24 h and imaged by IncuCyte FLR (Essen).

### Membrane leakage assay

Plasma membrane disruption during cell death was measured by lactate dehydrogenase (LDH) toxicity assay. PKLCs were seeded into 96-well plate, then treated with DMSO or the indicated compounds for different periods of time. Extracellular LDH activity was determined using LDH Release Assay Kit (Beyotime) according to the manufacturer’s instructions.

### Whole cell lysate preparation for immunoblotting analysis

Cells were collected from the 6-well plates or 6-cm petri dishes using a cell scraper, and then washed with PBS at room temperature. Cell pellets were resuspended in lysis buffer (20 mM Tris-HCl pH 7.4, 150 mM NaCl, 1% TritonX-100 and 5% glycerol), supplemented with protease inhibitors and phosphatase inhibitors (Share-bio), and incubated on ice for 30 min. After centrifugation at 13, 000×g for 10 min, total protein concentration of the supernatant was measured using Bradford Protein Assay Kit (Sangon Biotech). A corresponding dose of loading buffer (Solarbio) and lysis buffer was added to the sample to a final protein concentration of 2 mg/mL. The supernatants were boiled at 100 ℃ for 10 min and whole cell lysates of 20 - 40 μg proteins for each sample were subjected to SDS-PAGE for immunoblotting analysis.

### Isolation of TritonX-100-soluble and TritonX-100-insoluble fraction

Cells or were lysed with TritonX-100-containing lysis buffer as mentioned above. After centrifugation at 13,000 × g for 10 min, the supernatant was collected as TritonX-100-soluble fraction. The pellets were washed with PBS and resuspended in lysis buffer using high-speed tissue homogenizer. The suspension was added with loading buffer and lysis buffer to a final protein concentration of 2 mg/mL, and boiled at 100 ℃ for 10 min. Samples were saved as TritonX-100-insoluble fraction. For tumor tissue, freshly isolated tumors were lysed with TritonX-100-containing lysis buffer as mentioned above. Large tissue masses were then removed and tissue homogenates were centrifuged at 13,000 × g for 10 min. The supernatant was collected as TritonX-100-soluble fraction, Pellets were washed with PBS and resuspended in lysis buffer with 6 M urea for 6 h at 4℃. After centrifugation at 13, 000 × g for 10 min, the supernatant was collected as TritonX-100-insoluble, 6 M urea-soluble fraction.

### Plasma membrane isolation

Cells were collected, washed with PBS and resuspended in sucrose-containing buffer (10 mM Tris-HCl, pH 7.4, 250 mM sucrose, 5 mM MgCl_2_). After a 20 min-ice bath, the cells were disrupted with a Dounce homogenizer. Cell homogenates were collected and centrifuged at 17,000 × g for 15 min to remove nuclei and cellular debris. The supernatant was then re-centrifuged at 120,000 × g at 4 °C for 70 min. The pellet was collected as plasma membrane fraction.

### Immunoprecipitation

Whole cell lysates or indicated SEC fractions were incubated with anti-Flag or -V5 agarose beads at 4℃ overnight. Immunocomplexes were washed three times with lysis buffer and separated by SDS-PAGE for immunoblotting analysis using indicated antibodies.

### Size exclusion chromatography assay

Size exclusion chromatography assay was performed on an AKTA Purifier protein purification system using a superose 6 increase column according to the manufacturer’s instructions. In brief, 5 mg of total protein lysates were separated at a flow rate of 0.5 mL/min and 1 mL fractions collected at 4 ℃ in PBS with protease inhibitor cocktail and 1% TritonX-100. The molecular composition of each fraction was analyzed by immunoblotting using antibodies as indicated.

### Crystal violet assay

PKLC cells were seeded into 6-well plates and allowed to proliferate to 50% confluence. After grown in complete, glucose-deprived or glutamine-deprived DMEM under normoxia or hypoxia for different periods of times, the cells were stained with 0.5% crystal violet in 30% methanol at room temperature for 10 min, then washed with ddH_2_O. Air-dried plates were observed and imaged using a BZ-X810 microscope (KEYENCE).

### Lentivirus Production and Infection

HEK293T cells were co-transfected with lentiviral vectors carrying desired cDNAs or shRNAs and packaging plasmids (pMD2.G and psPAX2) by polyethylenimine (PEI) transfection reagent in serum-free medium. 8 h after transfection, the medium was changed to DMEM supplemented with 10% FBS and the virus-containing medium was collected 48 h later. For stable cell lines, target cells were seeded in 6-well plate and infected using indicated lentivirus in the presence of 10 µg/ml polybrene. 12 h after infection, the medium was changed. Gene and protein expression were analyzed 72 h post-infection by qRT-PCR and immunoblotting, respectively. For CRISPR-Cas9 mediated gene knockout, sgRNAs were expressed using a lentiviral vector LentiCRISPRv2. For RNA interference, shRNAs were expressed using a lentiviral vector pLKO.1-EGFP-Puro. For expression of exogenous proteins, wild-type, truncated, or mutant proteins were produced using eukaryotic expression vectors pHAGE-Puro or pLVX-Tet-On.

### shRNA mediated knock-down

Cells were infected with shRNA expressing lentiviral vectors as mentioned above. All the shRNAs targeting sequences for each gene are listed as follows: *Ripk1* (CGTGACTTTCACATTAAGATA), *Casp3* (CTCACGAAAGAACTGTACTTT), *Casp8* (CCTCCATCTATGACCTGACAT), *Gsdmd* (GATTGATGAGGAGGAATTAAT, GATGTCGTCGATGGGAACATT). A non-targeting shRNA construct (target sequence: TTCTCCGAACGTGTCACGTTT) served as negative control.

### CRISPR-Cas9 mediated gene knockout

For transfection-based delivery of CRISPR-Cas9 system, sgRNAs were cloned into pX330-mCherry. Cells were co-transfected with two sgRNA-expressing vectors by PEI reagent according to the manufacturer’s instructions. 36 h post transfection, mCherry-positive live cells were sorted by the FACSAria III Cell Sorter (BD Biosciences) and seeded in 10-cm petri dishes (100 cells per dish). After 10 days, single cell clones were picked up by trypsin digestion with cloning cylinders. Immunoblotting was used to screen for the KO clones. Knockout efficiency was confirmed by immunoblotting analysis. sgRNAs sequences are as follows: mouse *Mlkl* (GCACACGGTTTCCTAGACGC, CGCTAATTTGCAACTGCATC), mouse *Ripk1* (AGAAGAAGGGAACTATTCGC), mouse *Ripk3* (GAGGGTTCGGAGTCGTGTTC, ACCCTCCCTGAAACGTGGAC), mouse *Gsdmd* (CAGAGGCGATCTCATTCCGG, AGAAGGGAAAATTTCTGGTG). For lentiviral particle mediated CRISPR-Cas9 delivery, Cells were infected with sgRNA expressing lentiviral vectors as mentioned above. All the sgRNAs targeting sequences for each gene are listed as follows: mouse *Fadd* (TAGATCGTGTCGGCGCAGCG, TTCGTTTGCTCACGCGCTCG), mouse *Casp8* (CAAGAAGCAGGAGACCATCG, GTGGGATGTAGTCCAAGCAC, TGAGATCCCCAAATGTAAGC). A non-targeting sgRNA construct (target sequence: GGCCACGAGTTCGAGATCGA) served as negative control.

### Immunofluorescence staining and imaging

To detect protein localization by immunofluorescence in fixed PKLC cells, cells were first seeded on high-performance No.1.5 18 × 18 mm glass coverslips and allowed to grow overnight, followed by the indicated treatment. For immunofluorescence staining, cells were fixed with 4% PFA for 15 minutes, followed by permeabilization with 0.5% Triton X-100 for 5 minutes. The cells were then blocked with 2% BSA for 1 hour at room temperature. Primary antibodies were diluted in 2% BSA (mFlag 1:200, V5 1:200, RIP1 1:100, hFlag 1:200, Cleaved Caspase-8 1:100, Cleaved Caspase-3 1:400, phospho-MLKL (Ser345) 1:100, Flag-DyLight™ 488 1:100, V5-DyLight™ 650 1:100) and incubated for 1 hour at room temperature. After washing with 1 × DPBS 3 times, fluorescent secondary antibodies were diluted 1:1,000 in 2% BSA and incubated for 1 hour at room temperature. Samples were mounted in VECTASHIELD antifade mounting medium (Vector Lab).

Super-resolution images of PKLC-RIPK3^2×Fv^ cells stained with the antibodies against Flag-tag, V5-tag and RIPK1 were captured on SpinSR10 spinning disk confocal super resolution microscope (Olympus), the raw data were deconvoluted by cellSens using the enhanced ratio method. Other immunofluorescent images were captured on Dragonfly microscope (Andor) and raw data were deconvoluted by Fusion using the enhanced ratio method.

### PANoptosis-conditioned medium collection

PKLC-RIPK3^2×Fv^ cells were cultured in DMEM with 10% FBS and 1% penicillin-streptomycin. When the cells reach around 90 % confluence, cells were washed three times with PBS and changed into serum-free DMEM with 25 nM of B/B homodimerizer. The PKLC-RIPK3^2×Fv^ culture medium was harvested 8 h later after dimerizer treatment. The collected medium was successively centrifuged at 500 × g for 10 min, and 2,000 × g for 20 min. For Luminex cytokine analysis, PCM was supplemented with 0.1% TritonX-100 and protease inhibitors, incubated on ice for 30 min, and then passed through a 0.22 μm pore size filter.

### RNA isolation and real-time reverse transcription PCR analysis

Total RNA from cells was extracted by Trizol reagent (Life technology) according to the manufacturer’s instructions. cDNA was synthesized from total RNA by HiScript III All-in-one RT SuperMix (Vazyme) using poly (dT) primer, then diluted and used for real-time PCR with gene specific primers (5′-3′): mouse *Actb* (forward, GGCTGTATTCCCCTCCATCG; reverse, CCAGTTGGTAACAATGCCATGT); mouse *Ripk3* (forward, TCTGTCAAGTTATGGCCTACTGG; reverse, GGAACACGACTCCGAACCC); mouse *Ripk1* (forward, GAAGACAGACCTAGACAGCGG; reverse, CCAGTAGCTTCACCACTCGAC); mouse *Mlkl* (forward, AATTGTACTCTGGGAAATTGCCA; reverse, TCTCCAAGATTCCGTCCACAG). Real-time PCR was performed on an QuantStudio Dx Real-Time PCR System (Applied Biosystems) using ChamQ SYBR Color Master Mix (Vazyme).

### Luminex assay

The concentration of cytokines and chemokines in control CM and PCM were analyzed by a Luminex immunoassay following the manufacturer’s instructions (LXR-MultiDTM-23, −31; LabEx). Assays were performed on a Luminex 200 multiplexing instrument (Merck Millipore).

### Transwell chamber assay

iBMDM cells (0.8×10^6^ cells/insert) were seeded into Corning® Costar® Transwell inserts (8 μm pore size) suspended above 600 μL control CM or PCM. Following 24-hour incubation at 37°C/5% CO_2_, migrated cells on the bottom side of each well were imaged by DMi8 Inverted Microscope and counted. Four independent wells were analyzed.

### Statistical analysis

GraphPad Prism 10.0 software was used for data analysis. Data are presented as mean ± SD or SEM. Statistical significance was determined by t-tests (two-tailed) for two groups. P values less than 0.05 were considered statistically significant where, *p < 0.05, **p < 0.01, and ***p < 0.001.

## Data availability

All data and materials to draw the conclusions in this paper are presented in the main text, figures and the extended data figures. Further data can be received from the corresponding author on reasonable request. Source data are provided with this paper.

## Acknowledgments

We thank D. X., Y. L., H. P., S. Q., and L. C. for reagents. We thank S. H., M. Z., X. Y., B. H., and H. W. for helpful discussion. We thank X. D. for identification of GSDMD-deficient clones. We thank Chemical Biology Core Facility at SIBCB for technical assistance in time-lapsed imaging. This study is supported by the grants from the National Natural Science Foundation of China (82072573, 82273335 and 8240113215), Shanghai Municipal Health Commission (2022LJ015, 2022YQ039 and 20224Y0186).

## Author contributions

Y. Y., R. Z. and H. Z. conceptualized the study; Y. Y., Y. W^1^. and E. W. designed the methodology; Y. Y. and Y. W^1^. performed the majority of experiments and data analysis; Y. W^2^. performed the confocal microscopy and image analysis; L. S. did the mass spectrometry analysis; H. C. performed the bioinformatics analysis; Y. Y. wrote the manuscript with input from all authors; other authors offered technical inputs and discussed the results. Y. Z. and H. Z. acquired the fundings.

