## Supplementary Figures for "RIPK3-dependent sequential recruitment of MLKL and RIPK1 drives PANoptotic cell death and chemokine production"

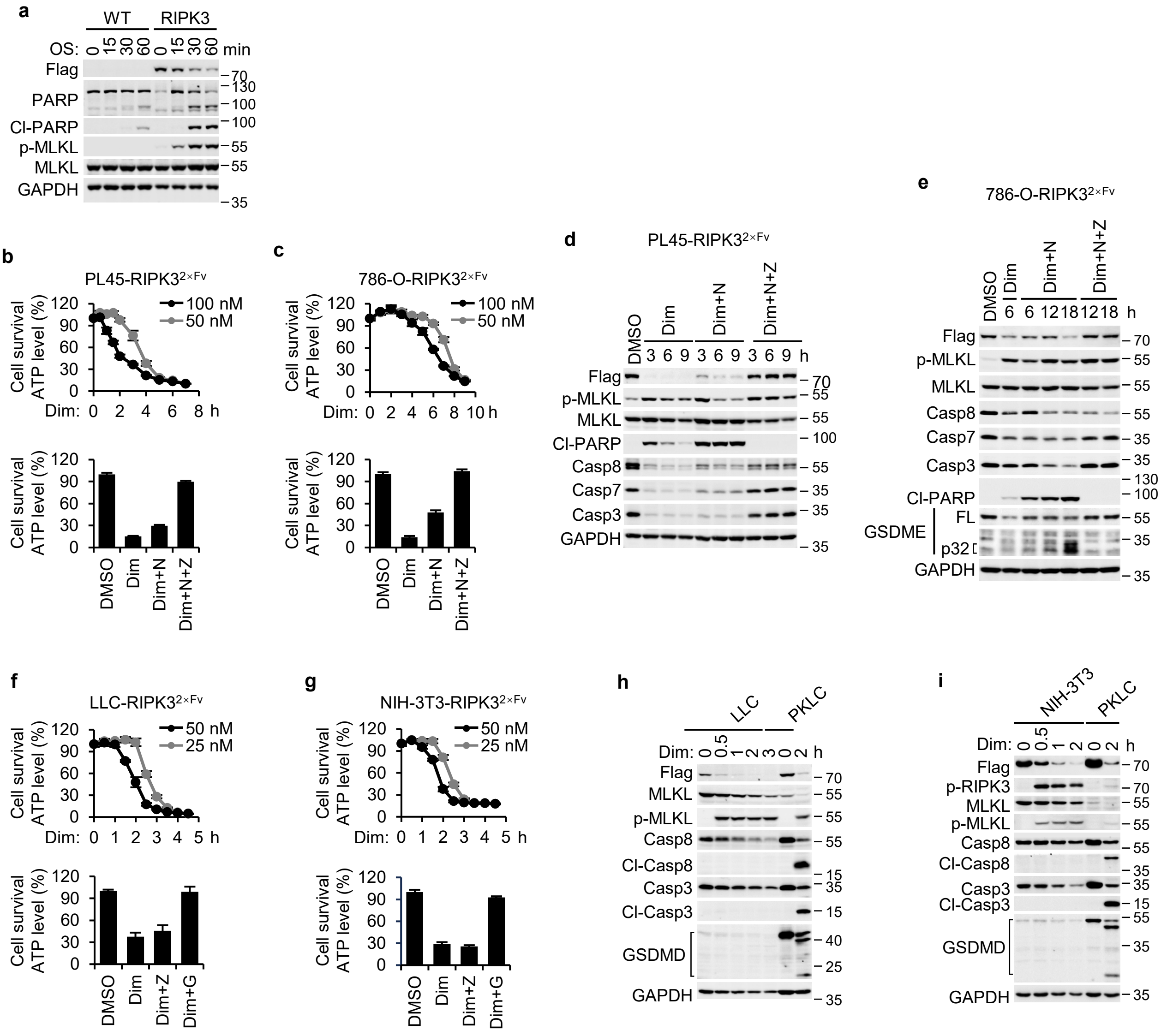


**Supplementary Fig. 1 | RIPK3 coordinates PANoptosis activation through cell type-specific mechanisms.**

**Supplementary Fig. 1 | RIPK3 coordinates PANoptosis activation through cell type-specific mechanisms. a** WT or RIPK3$^{2×Fv}$-restored PL45 cells were treated with OS for the indicated durations. Levels of p-MLKL (necroptosis marker), cleaved PARP (Cl-PARP, apoptosis marker) in cell lysates were detected by immunoblotting. GAPDH served as the loading control. Blots are representative of three independent experiments. **b, c** PL45-RIPK3$^{2×Fv}$, 786-O-RIPK3$^{2×Fv}$ cells were treated with 50 nM or 100 nM Dim for the indicated durations (upper panels in **b, c**), or with Dim, Dim + necrosulfonamide (NSA, N), Dim + N + Z-VAD (Z) for 9 hours (lower panels in **b, c**). Cell viability was determined by measuring ATP levels using the CellTiter-Glo kit. Data are presented as means ± SD of triplicate wells. **d, e** Whole-cell lysates from Dim-treated PL45-RIPK3$^{2×Fv}$ (**d**), 786-O-RIPK3$^{2×Fv}$ (**e**) cells, with or without indicated inhibitors, were analyzed by immunoblotting using the specified antibodies. Results are representative of three independent experiments. **f, g** LLC-RIPK3$^{2×Fv}$, NIH-3T3-RIPK3$^{2×Fv}$ cells were treated with 25 nM or 50 nM Dim for the indicated durations (upper panels in **f, g**), or with Dim, Dim + Z, Dim + G for 3 hours (lower panels in f, g). Cell viability was determined by measuring ATP levels using the CellTiter-Glo kit. Data are presented as means ± SD of triplicate wells. **h, i** Whole-cell lysates from Dim-treated LLC-RIPK3$^{2×Fv}$ (**h**), NIH-3T3-RIPK3$^{2×Fv}$ (**i**) cells, with or without indicated inhibitors, were analyzed by immunoblotting using the specified antibodies. Whole-cell lysates from Dim-treated PKLC-RIPK3$^{2×Fv}$ cells served as a positive control. Results are representative of three independent experiments.

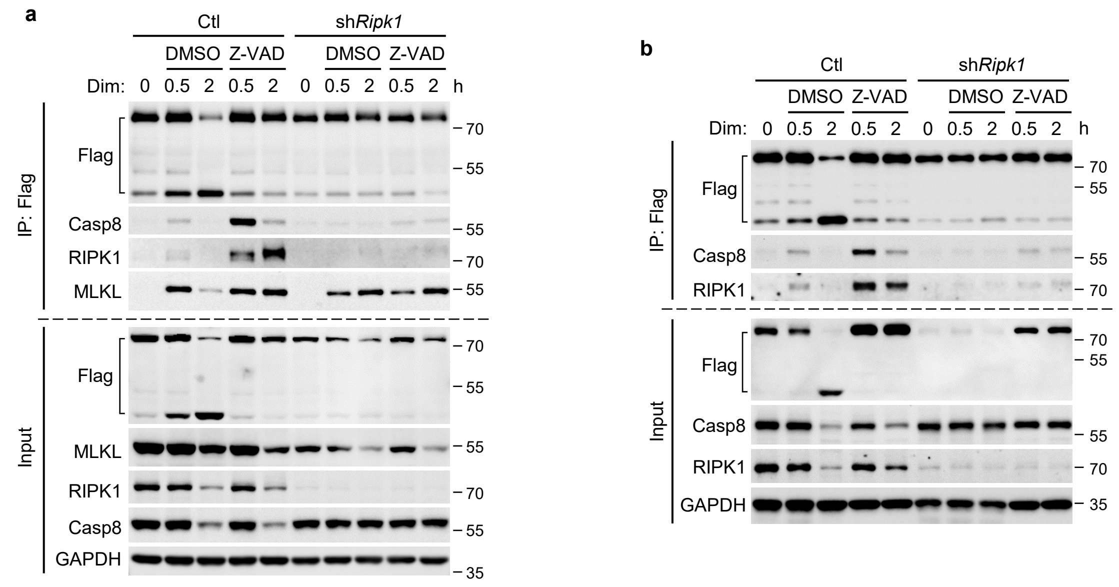


**Supplementary Fig. 2 | RIPK1 scaffolds RIPK3–Caspase-8 complex formation during RIPK3-PANoptosis.**

**Supplementary Fig. 2 | RIPK1 scaffolds RIPK3–Caspase-8 complex formation during RIPK3-PANoptosis.** WT and RIPK1-deficient PKLC-RIPK3$^{2\times Fv}$ (**a**) or MLKL-deficient PKLC-RIPK3$^{2\times Fv}$ (**b**) cells were treated with Dim or Dim + Z-VAD for the indicated durations. RIPK3 was immunoprecipitated from whole-cell lysates using anti-Flag resin. Lysates and immunocomplexes were analyzed by immunoblotting using antibodies against Flag, RIPK1, MLKL, and Casp8 as indicated. Results are representative of three independent experiments.

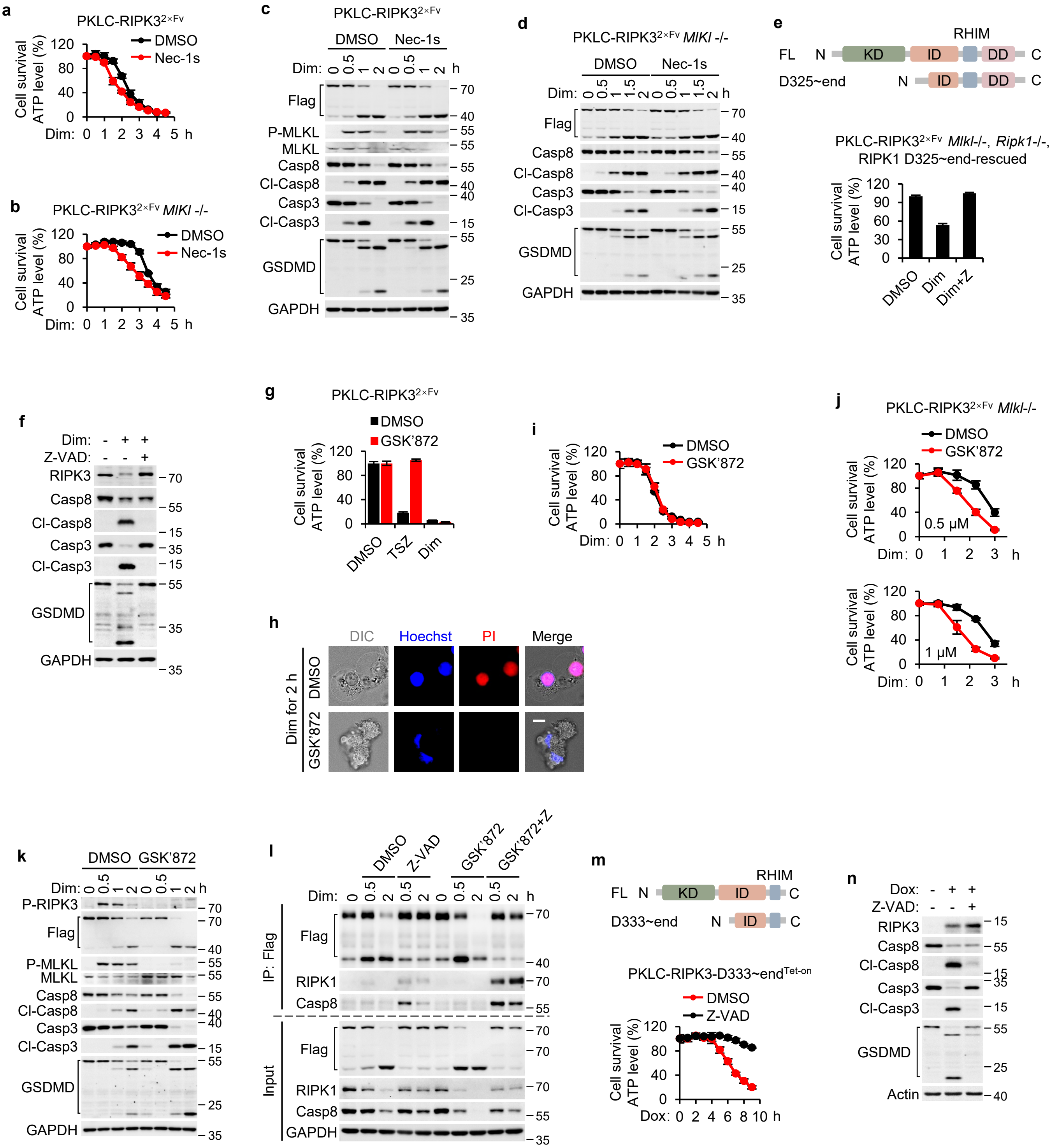


**Supplementary Fig. 3 | RIPK scaffolding mediates caspase activation during RIPK3-PANoptosis.**

**Supplementary Fig. 3 | RIPK scaffolding mediates caspase activation during RIPK3-PANoptosis. a-d** WT (**a, c**) or RIPK1-deficient (**b, d**) PKLC-RIPK3$^{2\times Fv}$ cells were treated with Dim or Dim + Nec-1s for the indicated durations. Cell viability was determined by measuring ATP levels using the CellTiter-Glo kit (**a, b**). Levels of p-MLKL (**c**), cleaved Caspase-8 (Cl-Casp8), cleaved Caspase-3 (Cl-Casp3), and cleaved GSDMD (Cl-GSDMD) (**c, d**) were detected by immunoblotting. **e, f** RIPK1$^{D325\sim end}$-rescued, MLKL-deficient PKLC-RIPK3$^{2\times Fv}$ cells were treated with Dim or Dim + Z-VAD (Z) for 4 hours. Cell viability was determined by measuring ATP levels using the CellTiter-Glo kit (**e**). Whole-cell lysates were analyzed by immunoblotting using the specified antibodies (**f**). **g-i** PKLC-RIPK3$^{2\times Fv}$ cells were treated with TSZ or Dim for 6 hours (**g**), or with Dim or Dim + GSK'872 for the indicated durations (**h, i**). Cell viability was determined by measuring ATP levels using the CellTiter-Glo kit (**g, i**). Cell morphology was visualized by wide-field light microscopy, and plasma membrane integrity was monitored by propidium iodide (PI) uptake (**i**). Nuclei were labeled with Hoechst. Scale bar, 10 µm. **j** MLKL-deficient PKLC-RIPK3$^{2\times Fv}$ cells were treated with DMSO or Dim (with or without GSK'872, 0.5 µM, upper panel; 1 µM, lower panel) for the indicated durations. Cell viability was determined by measuring ATP levels using the CellTiter-Glo kit. **k** PKLC-RIPK3$^{2\times Fv}$ cells were treated with Dim or Dim + GSK'872 for the indicated durations. Whole-cell lysates were analyzed by immunoblotting using the specified antibodies. **l** MLKL-deficient PKLC-RIPK3$^{2\times Fv}$ cells were treated with Dim or Dim + GSK'872 for the indicated durations. RIPK3 was immunoprecipitated from whole-cell lysates using anti-Flag resin. Lysates and immunocomplexes were analyzed by immunoblotting using antibodies against Flag, RIPK1, and Casp8. **m, n** Dox-inducible RIPK3$^{D333\sim end}$ truncation-expressing PKLCs were treated with Dox or Dox + Z for the indicated durations. Cell viability was determined by measuring ATP levels using the CellTiter-Glo kit (**m**). Whole-cell lysates were analyzed by immunoblotting using the specified antibodies (**n**). Data are presented as means ± SD of triplicate wells (**a, b, e, g, i, j, m**). Results are representative of three independent experiments (**c, d, f, k, l, n**).

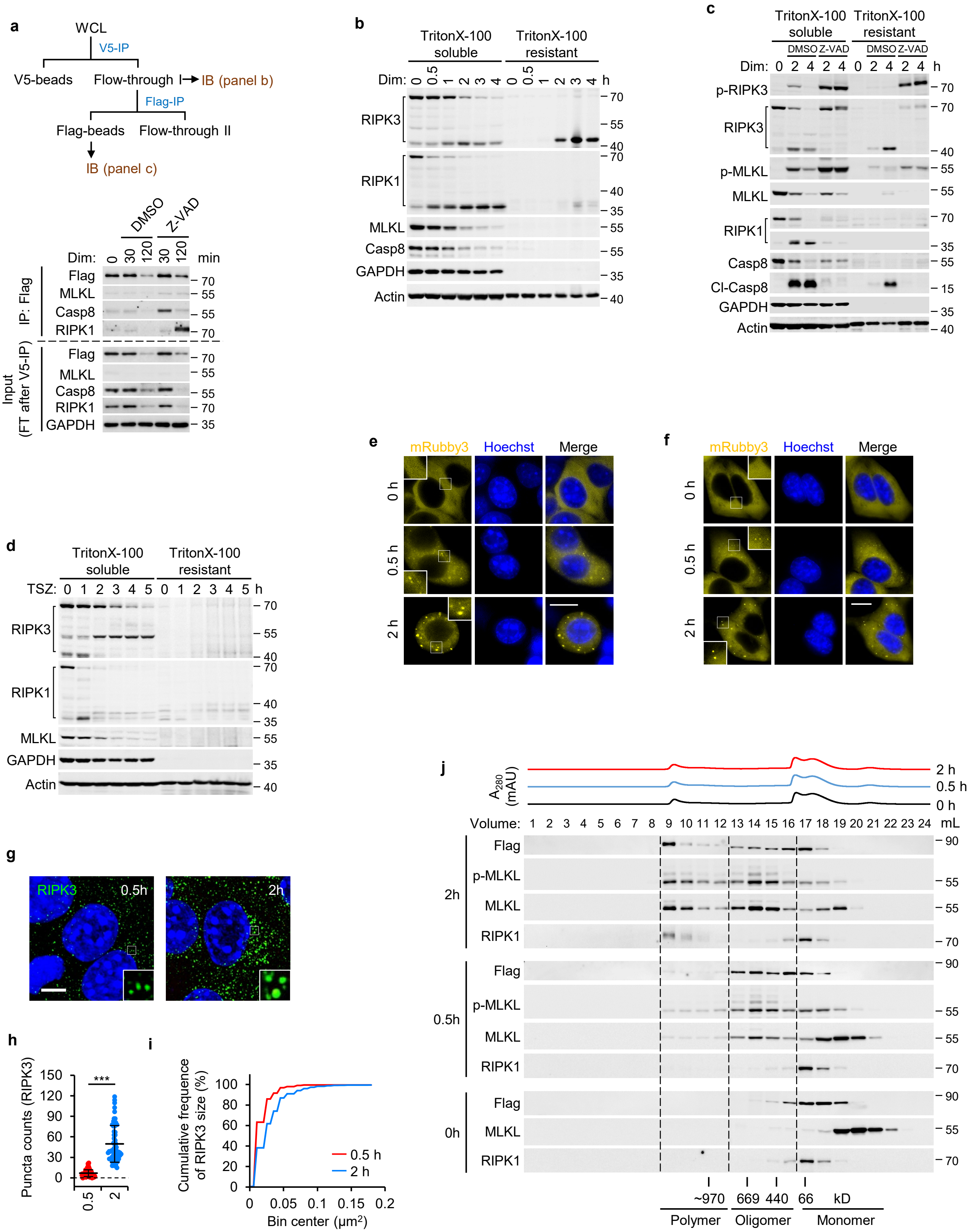


**Supplementary Fig. 4 | RIPK3 hierarchically assembles multimeric scaffold architectures to drive PANoptosis execution.**

**Supplementary Fig. 4 | RIPK3 hierarchically assembles multimeric scaffold architectures to drive PANoptosis execution. a** The MLKL-removed lysates (flow-through I, FT) after V5 immunoprecipitation (IP) in Fig. 4**d** were immunoprecipitated with anti-Flag resin. Immunocomplexes were analyzed by immunoblotting using the indicated antibodies. **b-d** PKLC-RIPK3$^{2\times Fv}$ cells were treated with Dim (**b, c**), Dim + Z-VAD (**c**), or TSZ (**d**) for the indicated durations. The localization of RIPK3, RIPK1, MLKL, and Casp8 in Triton X-100-soluble and Triton X-100-resistant fractions was analyzed by immunoblotting using the indicated antibodies. **e, f** Representative live-cell images of WT MLKL- (**e**) and N-terminally V5-tagged MLKL- (**f**) expressing PKLC-RIPK3$^{2\times Fv}$ cells treated with Dim for the indicated durations. RIPK3 was traced by fusing an mRuby3 fluorescent protein at its C-terminus. **g** Representative immunofluorescence images of PKLC-RIPK3$^{2\times Fv}$ cells stained with anti-Flag antibody after treatment with Dim + Z for the indicated durations. Scale bar, 10 μm. **h** Quantification of RIPK3 puncta counts based on the fluorescence images in **g**. Data are presented as means ± SD. Statistical significance was determined by two-sided unpaired Student's t-tests. ***$p < 0.001$. **i** Cumulative distribution plots depicting RIPK3 puncta size in PKLC-RIPK3$^{2\times Fv}$ cells treated with Dim for the indicated durations. **j** Triton X-100-soluble lysates from Dim + Z-treated PKLC-RIPK3$^{2\times Fv}$ cells were fractionated using a Superose 6 Increase size-exclusion chromatography column. Collected fractions were analyzed by immunoblotting using antibodies against RIPK3, RIPK1, MLKL, and p-MLKL. UV absorbance curves from size-exclusion chromatography (SEC) analysis are displayed at the top.

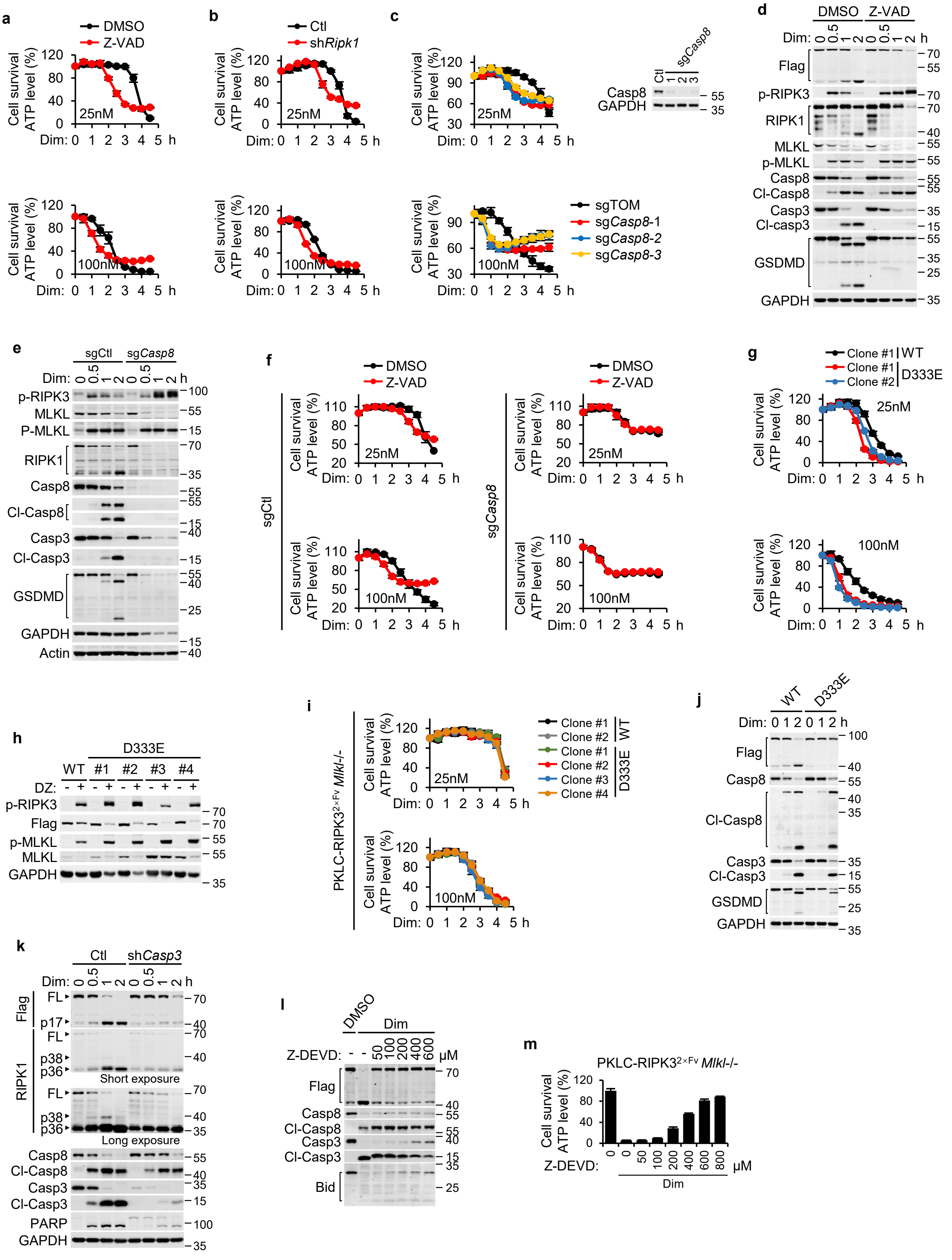

**Supplementary Fig. 5 | Caspase-mediated RIPK3 cleavage suppresses PANoptosis by inhibiting MLKL-dependent membrane permeabilization.**

**Supplementary Fig. 5 | Caspase-mediated RIPK3 cleavage suppresses PANoptosis by inhibiting MLKL-dependent membrane permeabilization. a-f** Wild-type (**a, d**), RIPK1-deficient (**b**), or caspase-8-deficient (**c, e, f**) PKLC-RIPK3$^{2\times Fv}$ cells were treated with Dim (**a-f**) or Dim + Z-VAD (**a, d, f**) for the indicated durations. **g-j** WT (**g, h**) or MLKL-deficient (**i, j**) PKLC cells expressing WT RIPK3$^{2\times Fv}$ or its caspase-resistant mutant D333E$^{2\times Fv}$ were treated with Dim (25 nM, upper panel; 100 nM, lower panel) for the indicated durations. Cell viability was determined by measuring ATP levels using the CellTiter-Glo kit (**a-c, f, g, i**). Whole-cell lysates were analyzed by immunoblotting using the specified antibodies (**d, e, h, j**). **k** Cleavage of RIPK3 and RIPK1 in wild-type or caspase-3-deficient PKLC-RIPK3$^{2\times Fv}$ cells treated with Dim for the indicated durations was detected by immunoblotting using the specified antibodies. **l, m** MLKL-deficient PKLC-RIPK3$^{2\times Fv}$ cells pretreated with Z-DEVD-FMK (Z-DEVD), a caspase-3/7 inhibitor, at the indicated concentrations were treated with Dim for 4 hours. Cell viability was determined by measuring ATP levels using the CellTiter-Glo kit (**m**). Cleavage of RIPK3, caspase-8, and caspase-3 was detected by immunoblotting using the specified antibodies (**l**). Data are presented as means ± SD of triplicate wells (**a-c, f, g, i, m**). Results are representative of three independent experiments (**d, e, h, j, k, l**).

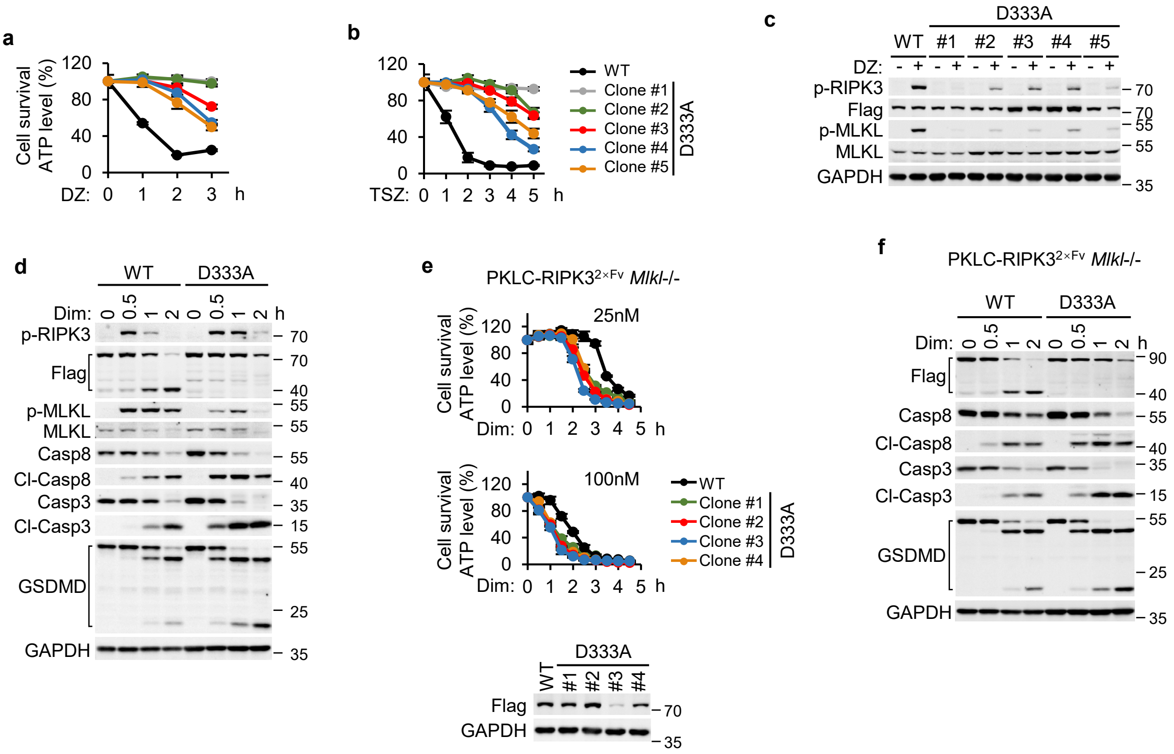


**Supplementary Fig. 6 | Caspase-cleavage-resistant RIPK3-D333A mutant suppresses necroptosis through kinase activity restriction.**

**Supplementary Fig. 6 | Caspase-cleavage-resistant RIPK3-D333A mutant suppresses necroptosis through kinase activity restriction. a-c** WT RIPK3$^{2\times Fv}$-expressing PKLC or various RIPK3-D333A$^{2\times Fv}$ mutant-expressing PKLC clones were treated with DZ (**a, c**) or TSZ (**b**) for the indicated durations. Cell viability was determined by measuring ATP levels using the CellTiter-Glo kit (**a, b**). Phosphorylation of RIPK3 and MLKL in WT RIPK3$^{2\times Fv}$-expressing PKLC or D333A$^{2\times Fv}$ mutant-expressing PKLC clones after 4 hours of DZ treatment was detected by immunoblotting using the specified antibodies (**c**). **d-f** PKLC (**d**) or MLKL-deficient PKLC (**e, f**) cells expressing WT RIPK3$^{2\times Fv}$ or RIPK3-D333A$^{2\times Fv}$ mutants were treated with Dim for the indicated durations. Cell viability was determined by measuring ATP levels using the CellTiter-Glo kit (**e**). Whole-cell lysates were analyzed by immunoblotting using the specified antibodies (**d, f**). Data are presented as means ± SD of triplicate wells (**a, b, e**). Results are representative of three independent experiments (**c, d, f**).

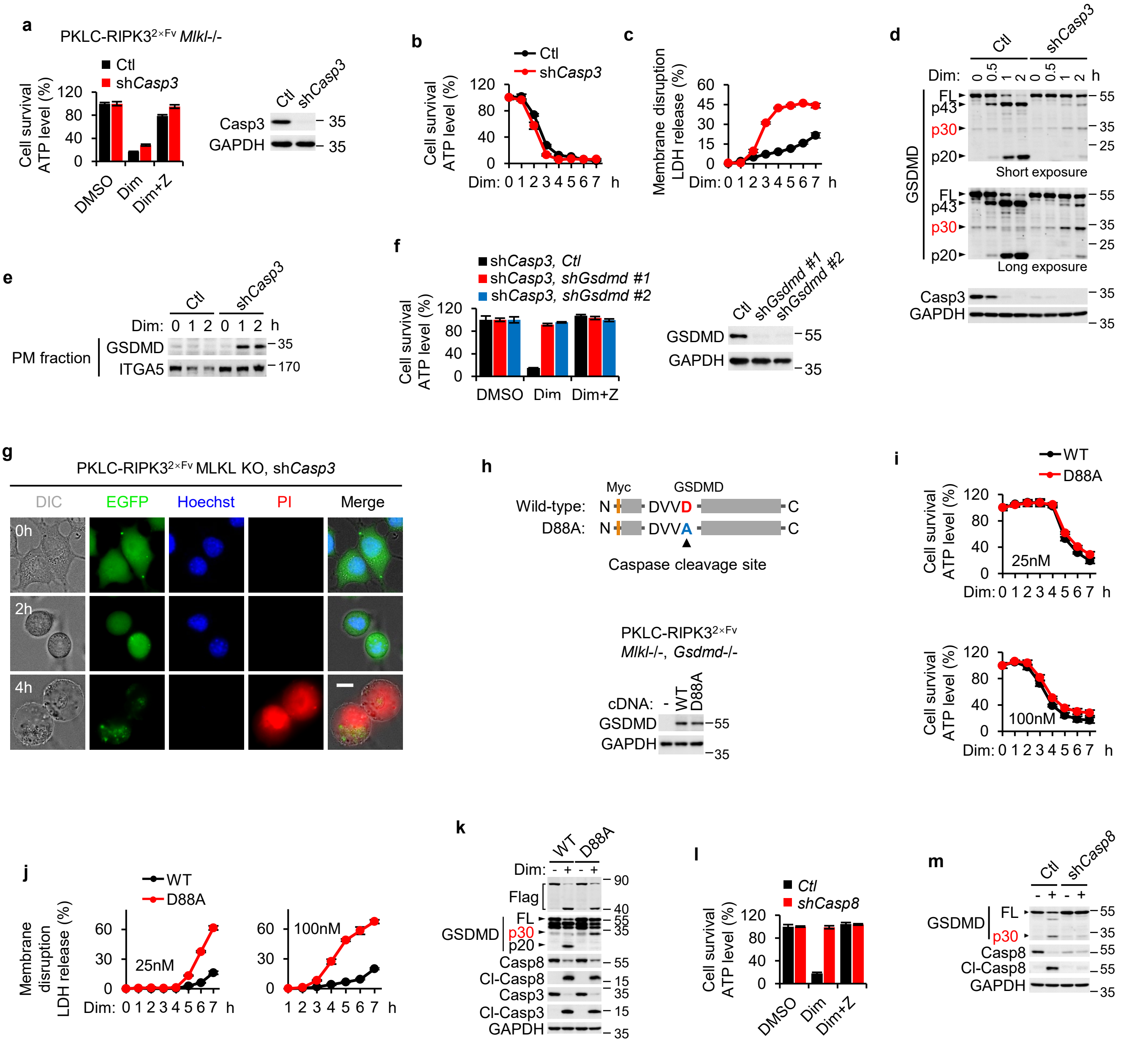

**Supplementary Fig. 7 | Caspase-3-dependent cleavage of GSDMD at Asp88 suppresses pyroptotic membrane permeabilization.**

**Supplementary Fig. 7 | Caspase-3-dependent cleavage of GSDMD at Asp88 suppresses pyroptotic membrane permeabilization. a-e** WT or caspase-3-deficient PKLC-RIPK3$^{2\times Fv}$ cells (with MLKL deficiency) were treated with Dim or Dim + Z for 4 hours (**a**) or with Dim for the indicated durations (**b-e**). Cell viability was determined by measuring ATP levels using the CellTiter-Glo kit (**a, b**). Plasma membrane disruption was assessed by measuring lactate dehydrogenase (LDH) release into the cell medium (**c**). GSDMD cleavage (**d**) and plasma membrane translocation (**e**) were detected by immunoblotting using the specified antibodies. **f** GSDMD-knockdown PKLC-RIPK3$^{2\times Fv}$ cells with MLKL/caspase-3 double deficiency were treated with Dim or Dim + Z for 4 hours. Cell viability was determined by measuring ATP levels using the CellTiter-Glo kit. **g** Representative images of MLKL-/Casp3-deficient PKLC-RIPK3$^{2\times Fv}$ cells treated with Dim for the indicated durations. Cell morphology was visualized by wide-field light microscopy, and plasma membrane integrity was monitored by propidium iodide (PI) uptake and EGFP release. Nuclei were labeled with Hoechst. Scale bar, 10 μm. **h-k** WT or D88A (a caspase-3-resistant mutant) GSDMD-expressing PKLC-RIPK3$^{2\times Fv}$ cells (**h**) with MLKL deficiency were treated with Dim for the indicated durations. Cell viability was determined by measuring ATP levels using the CellTiter-Glo kit (**i**). Plasma membrane disruption was assessed by measuring LDH release into the cell medium (**j**). Cleavage of GSDMD and caspases was detected by immunoblotting using the specified antibodies (**k**). **l, m** WT and caspase-8-knockdown PKLC-RIPK3$^{2\times Fv}$ cells with MLKL/caspase-3 double deficiency were treated with Dim or Dim + Z for 4 hours. Cell viability was determined by measuring ATP levels using the CellTiter-Glo kit (**l**). GSDMD cleavage was detected by immunoblotting (**m**). Data are presented as means ± SD of triplicate wells (**a-c, f, i, j, l**). Results are representative of three independent experiments (**d, e, k, m**).

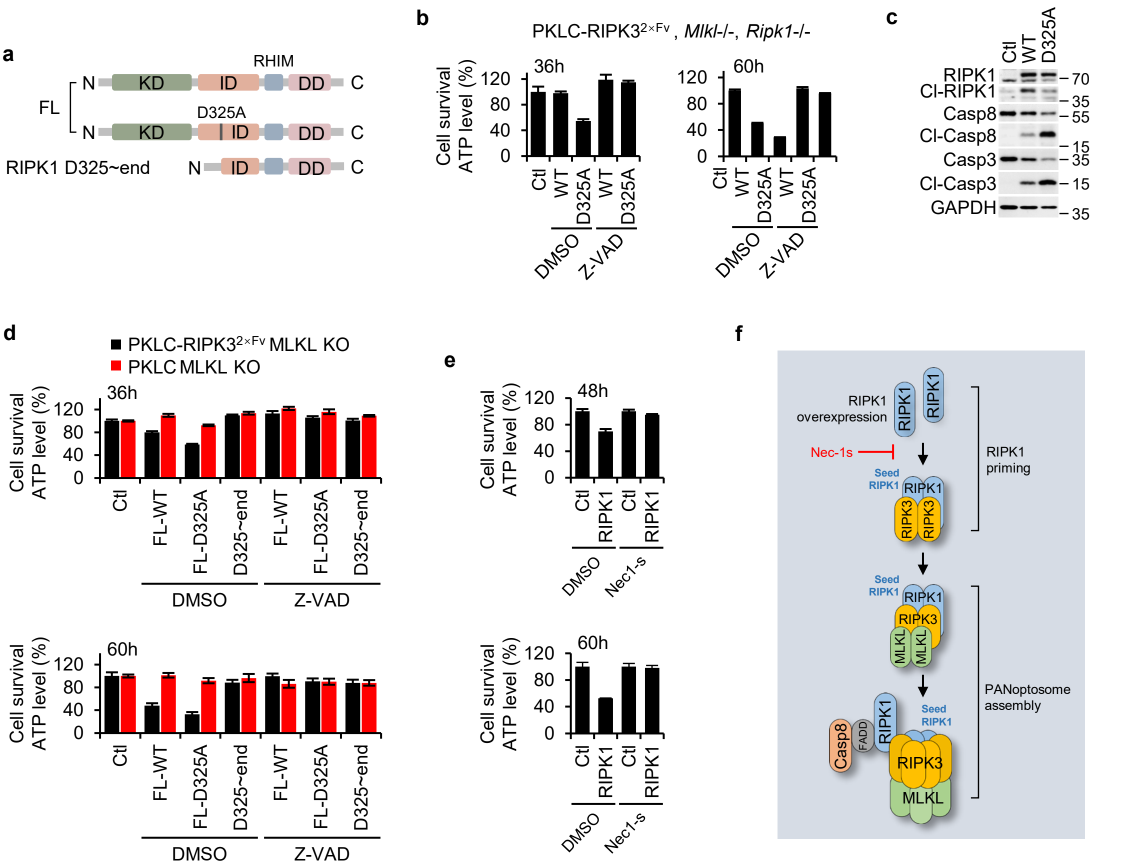


**Supplementary Fig. 8 | RIPK1 kinase-dependent priming licenses RIPK3 activation followed by kinase-independent recruitment executing caspase engagement.**

**Supplementary Fig. 8 | RIPK1 kinase-dependent priming licenses RIPK3 activation followed by kinase-independent recruitment executing caspase engagement. a** Schematic representation of WT RIPK1, RIPK1 D325A mutant, and C-terminal truncations of RIPK1. **b, c** MLKL- and RIPK1-deficient PKLC-RIPK3$^{2\times Fv}$ cells were infected with concentrated lentiviral particles encoding WT RIPK1 or RIPK1 D325A mutant for 36 hours (**b, c**) or 60 hours (**b**) as indicated. Cell viability was determined by measuring ATP levels using the CellTiter-Glo kit (**b**). Cleavage of caspase-8 and caspase-3 was detected by immunoblotting using the specified antibodies (**c**). Results are representative of three independent experiments. **d, e** MLKL-deficient PKLC-RIPK3$^{2\times Fv}$ (**d**) or PKLC (**d, e**) cells were infected with concentrated lentiviral particles encoding WT RIPK1 (**d, e**), RIPK1 D325A mutant, or RIPK1 D325~end truncation (**d**) for 36 hours, 60 hours (**d, e**), or 48 hours (**e**) as indicated. Cell viability was determined by measuring ATP levels using the CellTiter-Glo kit. Data are presented as means ± SD of triplicate wells (**b, d, e**). **f** Schematic illustrating RIPK1-initiated RIPK3-dependent PANoptosis.

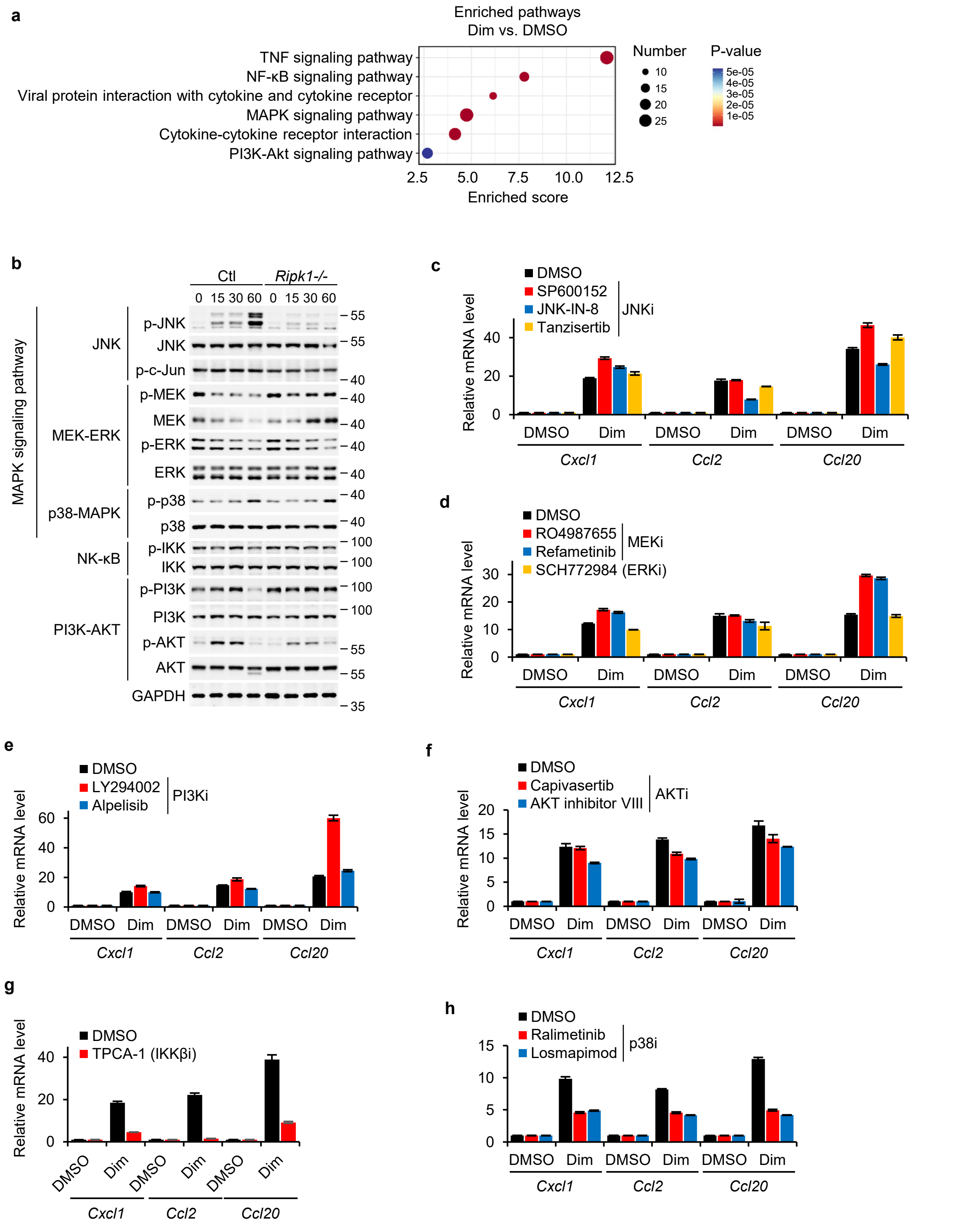


**Supplementary Fig. 9 | IKKβ mediates RIPK1-dependent NF-κB activation to promote chemokine production during RIPK3-PANoptosis.**

**Supplementary Fig. 9 | IKKβ mediates RIPK1-dependent NF-κB activation to promote chemokine production during RIPK3-PANoptosis. a** KEGG pathway enrichment analysis of differentially expressed genes (DEGs) in PKLC-RIPK3$^{2×Fv}$ cells treated with DMSO or Dim for 1 hour. The top six significantly enriched biological processes are shown ($P < 0.05$).
**b** Immunoblotting analysis of MAPK, NF-κB, and PI3K-AKT signaling pathway activation markers in MLKL-deficient PKLC-RIPK3$^{2×Fv}$ cells (WT or RIPK1-deficient) treated with Dim for the indicated durations. Results are representative of three independent experiments. **c-h** qRT-PCR quantification of *Cxcl1*, *Ccl2*, and *Ccl20* mRNA levels in MLKL-deficient PKLC-RIPK3$^{2×Fv}$ cells treated with 50 nM Dim ± indicated inhibitors of JNK (**c**), MEK/ERK (**d**), IKK (**e**), p38 (**f**), PI3K (**g**), or AKT (**h**) for 1 hour. Data are presented as means ± SD of triplicate wells.
